# Deep-learning Based Pathological Assessment of Frozen Procurement Kidney Biopsies Predicts Graft Loss and Guides Organ Utilization: A Large-scale Retrospective Study

**DOI:** 10.1101/2023.05.31.543098

**Authors:** Zhengzi Yi, Caixia Xi, Madhav C Menon, Paolo Cravedi, Fasika Tedla, Alan Soto, Zeguo Sun, Keyu Liu, Jason Zhang, Chengguo Wei, Man Chen, Wenlin Wang, Brandon Veremis, Monica Garcia-barros, Abhishek Kumar, Danielle Haakinson, Rachel Brody, Lorenzo Gallon, Philip O’Connell, Maarten Naesens, Ron Shapiro, Robert Colvin, Stephen Ward, Fadi Salem, Weijia Zhang

**Author notes:** Correspondence: Dr. Weijia Zhang Division of Nephrology Department of Medicine, Icahn School of Medicine at Mount Sinai One Gustave L Levy Place, Box 1243 New York, NY 10029, Dr. Fadi Salem, Department of Laboratory Medicine & Pathology Mayo Clinic, Rochester, MN 55902, Dr. Stephen Ward, Department of Pathology, Molecular and Cell based Medicine, Icahn School of Medicine at Mount Sinai, New York, NY 10029.

## Abstract

**Background:** Lesion scores on procurement donor biopsies are commonly used to guide organ utilization. However, frozen sections present challenges for histological scoring, leading to inter- and intra-observer variability and inappropriate discard.

**Methods:** We constructed deep-learning based models to recognize kidney tissue compartments in H&E stained sections from procurement biopsies performed at 583 hospitals nationwide in year 2011-2020. The models were trained and tested respectively on 11473 and 3986 images sliced from 100 slides. We then extracted whole-slide abnormality features from 2431 kidneys, and correlated with pathologists’ scores and transplant outcomes. Finally, a Kidney Donor Quality Score (KDQS) incorporating digital features and the Kidney Donor Profile Index (KDPI) was derived and used in combination with recipient demographic and peri-transplant characteristics to predict graft loss or assist organ utilization.

**Results:** Our model accurately identified 96% and 91% of normal/sclerotic glomeruli respectively; 94% of arteries/arterial intimal fibrosis regions; 90% of tubules. Three whole-slide features (Sclerotic Glomeruli%, Arterial Intimal Fibrosis%, and Interstitial Fibrosis%) demonstrated strong correlations with corresponding pathologists’ scores (n=2431), but had superior associations with post-transplant eGFR (n=2033) and graft loss (n=1560). The combination of KDQS and other factors predicted 1- and 4-year graft loss (discovery: n=520, validation: n=1040). Finally, by matching 398 discarded kidneys due to “biopsy findings” to transplanted population, the matched transplants from discarded KDQS<4 group (110/398, 27.6%) showed similar graft survival rate to unmatched transplanted kidneys (2-, 5-year survival rate: 97%, 86%). KDQS ≥ 7 (37/398, 9.3%) and 1-year survival model score ≥ 0.55 were determined to identify possible discards (PPV=0.92).

**Conclusion:** This deep-learning based approach provides automatic and reliable pathological assessment of procurement kidney biopsies, which could facilitate graft loss risk stratification and organ utilization.

**Translational Statement:** This deep-learning based approach provides rapid but more objective, sensitive and reliable assessment of deceased-donor kidneys before transplantation, and improves the prognostic value of procurement biopsies, thus could potentially reduce inappropriate discard and stratify patients needing monitoring or preventative measures after transplantation. The pipeline can be integrated into various types of scanners and conveniently generates report after slide scanning. Such report can be used in conjunction with pathologists’ report or independently for centers lacking renal pathologists.

## Introduction

Kidney transplantation prolong long-term survival of patients with end-stage renal disease waitlisted on dialysis[1]. In US, despite a 7-9% increase in the number of new candidates each year, only a quarter of waitlisted patients receive a deceased-donor kidney transplant within 5 years, and even fewer receive a living-donor transplant[2]. On the other hand, 20% of recovered organs are discarded annually (29% during the pandemic[2, 3]). This discard rate in US was reported to be nearly twice higher than that of France[4]. Among the reasons reported for discard, “Biopsy Findings” was the most common factor[5–7].

Procurement biopsies are performed during allocation of deceased donor kidneys to ascertain histological lesions including glomerulosclerosis, arteriosclerosis, and interstitial fibrosis[8]. However, conflicting results of the association of these histological findings and post-transplant graft outcomes[9–15] have diminished the utility of procurement biopsies in practice, as concerns are growing that these biopsies may lead to inappropriate discard. The major limitations of procurement biopsies are the use of frozen sections as the freezing artifacts could lead to pathological misinterpretations [16, 17], and the inter-rater variation among on-call pathologists and even among experts[18–20]. Therefore, Lentine et al.[20] proposed that the application of artificial intelligence (AI) algorithms could be one promising approach so as to allow robust histological assessment.

In recent years, deep-learning based AI techniques have been successfully applied to kidney pathology given their ability to extensively learn from large-scale image data[21–23]. Previous research mostly focused on glomerular disease classification only or post-transplant diagnosis[24], while few recent studies have shown good prediction of tissue compartments through pixel-level segmentation (U-net) or instance level detection (Mask R-CNN) method[25–28]. However, to our knowledge, none is reported to detect arteriosclerosis and be applied to large amount of frozen sections which are difficult to interpret by pathologists[18, 19, 29]. We aimed to utilize AI techniques to assist in the pathological evaluation of procurement biopsies, and eventually to improve graft loss risk estimation and potentially prevent unnecessary organ discard.

## Methods

### Study cohort and biopsy slides

The study included a total number of 2431 of deceased donor kidneys from 583 hospitals across United States partnered with the Organ Procurement and Transplantation Network (OPTN) (Figure1A). These kidneys were further discarded or allocated to 95 transplant centers nationwide (Table S1) for transplantation. The cohort consisted of 2033 transplanted kidneys in the years 2011-2016 (n=1560) and 2019-2020 (n=473). The kidneys transplanted in 2011-2016 were randomly divided into a discovery set (n=520) and a validation set (n=1040) for intermediate-term graft survival analyses. In addition, to explore organ utilization following latest standard, we first identified 1209 kidneys discarded in our cohort in recent five years (2015-2020), among which 398 out of 862 kidneys discarded due to biopsy findings with available slides were specifically selected for estimation of organs saved, i.e. organs that were discarded due to biopsy findings but potentially could have been transplanted successfully.

H&E-stained slides from frozen sections of needle procurement biopsies were used in this study. They were digitalized with the Aperio (slides in 2019-2020) and Hamamatsu (remaining slides) scanners at 20x resolution. Each H&E slide was scored by one transplant pathologist and reviewed by at least one independent pathologist at the time of biopsy; multiple pathologists were involved between 2011-2020 and three lesion scores: Sclerotic Glomeruli% (GS%), Arterial Intimal Fibrosis% (AIF%) Interstitial Fibrosis% (IF%) were recorded. Two public composite scores: Donor Chronic Damage Score[30] and Leuven Donor Risk Score[31] were evaluated in our cohort in order to compare to our digital composite score. Due to the unavailability of tubular atrophy score, Remuzzi Score[32, 33] was not evaluated here.

The demographic and clinical data were obtained from OPTN under institutional IRB approval (HS#: STUDY-21-01627). The OPTN data system includes data on all donors, wait-listed candidates, and transplant recipients in the US, submitted by the members of the OPTN. The Health Resources and Services Administration (HRSA), U.S. Department of Health and Human Services provide oversight to the activities of the OPTN contractor.

### Deep-learning based tissue compartment recognition and whole slide investigation

We first constructed tissue compartment recognition models, then built a pipeline transforming raw whole slide images (WSIs) into digital pathological reports (Figure 1B). In stage I, the compartments, including normal and sclerotic glomeruli, arteries and intimal fibrosis regions, and tubules were annotated from 100 large WSIs. The annotated sections were further pre-processed into 15459 fixed-sized image tiles (11473 as training set, 3986 as testing set) at 10x objective-power for model construction via 5-fold cross-validation. We used Mask R-CNN structure for compartment detection and U-net segmentation structure for tubule prediction enhancement. In Stage II, a pipeline was designed to take scanned WSI as input, and automatically extracted tissue sections which were then send to pre-trained deep learning models for compartments recognition, finally aggregated to estimate slide-level lesion scores and generated prediction images and reports as output. The whole WSI investigation process supports file formats from different types of scanner [34] and takes 30 minutes. Detailed methods were described in supplementary methods.

**Figure 1.**
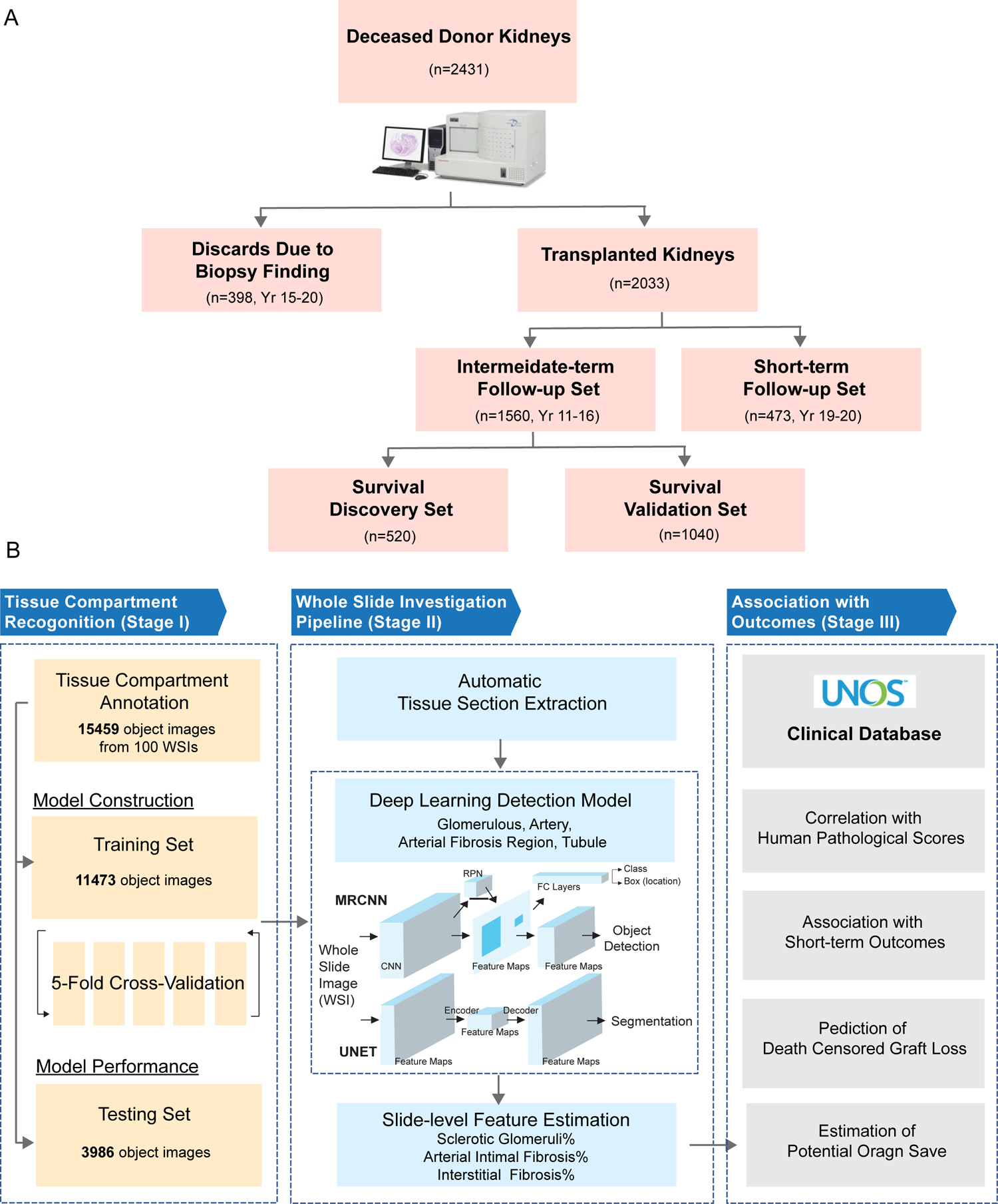
Study cohort and research workflow. **A)** Cohort description. We scanned and processed H&E stained slides of procurement biopsies from 2431 deceased donor kidneys which included 398 discarded kidneys due to biopsy findings (years 2015-2020) and 2033 transplanted kidneys (years 2011-2016 and years 2019-2020). The transplanted cohort contained 1560 kidneys transplanted in years 2011-2016 and 473 kidneys transplanted in years 2019-2020. All kidneys were used for correlation analyses with pathologist scores. All transplanted kidneys were used for association analyses with post-transplant short-term outcomes within 24-months while the kidneys transplanted in years 2011-2016 were used for survival analyses. To derive a composite score in assisting graft loss prediction and organ utilization, we further divided kidneys transplanted in years 2011-2016 cohort into a n=520 discovery set and a n=1040 validation set. **B)** The workflow consists of three stages. In the first stage (tissue compartment recognition), kidney tissue sections were annotated and were divided into training set (11473 images) and testing set (3986 images) for deep-learning tissue compartment recognition model construction. In the second stage (whole slide feature extraction), the pre-trained model was embedded into a whole slide investigation pipeline which accepted scanned raw image as input and generated whole side prediction image as well as slide level pathological abnormality report. In the third stage (clinical investigation), we examined associations of digital abnormality scores and pathologist scores with post-transplant graft outcomes.

### Statistical analysis

Correlations of digital scores with pathologist scores were assessed by Pearson’s correlation for Sclerotic Glomeruli% and by Spearman’s correlation for Arterial Intimal Fibrosis% and Interstitial Fibrosis%. When evaluating association of biopsy scores with graft outcomes or graft loss, multivariate linear regressions or cox-regressions were performed respectively by adjusting for recipient related factors and peri-transplant characteristics. Prediction of graft loss at specific time point was performed by logistic regression. The matching of discarded kidneys to transplanted kidneys was performed by the R package “MatchIt”[35]. Detailed methods and adjusting confounders were provided in supplementary methods.

## Results

### Cohort description

Donor and recipient characteristics are summarized in Table 1. 2033 transplant recipients with a mean age of 55.07±13.07 years received kidneys from 1489 donors with a mean age of 45.16±13.24 years and Kidney Donor Profile Index (KDPI)[36] of 0.55 ±0.25. Most donors were Caucasian (61.25%) while ethnic backgrounds in recipients were more diverse. The recipients were followed up for average of 4.4 years, with a graft loss rate of 15.64%. The discarded kidneys were from 294 donors with a mean average age of 50.78±11.88 and KDPI of 0.72±0.20.

**Table 1.** Demographic and clinical characteristics of recipients and donors in transplanted and discarded cohorts. Continuous variables are shown as mean value ± standard deviation, categorical variables are shown as count (percentage).

| **Recipient/Transplant Characteristics** | **Transplanted (n=2033 recipients)** | **Donor Characteristics** | **Transplanted (n=1489 donors)** | **Discarded (n=294 donors)** |
| --- | --- | --- | --- | --- |
| Recipient Age | 55.07 ±13.07 | Donor Age | 45.16 ±13.24 | 50.78±11.88 |
| Recipient Gender |  | Donor Gender |  |  |
| Female | 746(36.69) | Female | 598(40.16) | 129(43.88) |
| Male | 1287(63.31) | Male | 891(59.84) | 165(56.12) |
| Recipient Race |  | Donor Race |  |  |
| Asian | 224(11.02) | Asian | 38(2.55) | 3(1.02) |
| African American | 794(39.06) | African American | 228(15.31) | 58(19.73) |
| Hispanic | 502(24.69) | Hispanic | 296(19.88) | 38(12.93) |
| Caucasian | 499(24.55) | Caucasian | 912(61.25) | 189(64.29) |
| Other | 14(0.68) | Other | 15(1.01) | 6(2.04) |
| Recipient Pre-transplant Dialysis | | KDPI | 0.55 ±0.25 | 0.72±0.20 |
| No | 222(10.93) |  |  |  |
| Yes | 1808(88.98) |  |  |  |
| Unknown | 2(0.09) |  |  |  |
| Estimated Post-Transplant Survival (EPTS) score | 0.55±0.29 | | | |
| Pump Usage |  |  |  |  |
| No | 137(6.74) |  |  |  |
| Yes | 1896(93.26) |  |  |  |
| Cold Ischemia Time (hours) | 25.82 ±10.89 | | | |
| Induction Type |  |  |  |  |
| Lymphocyte Non-depletion | 398(19.58) |  |  |  |
| Lymphocyte Depletion | 1517(74.62) |  |  |  |
| None | 118(5.8) |  |  |  |
| Death Censored Graft Loss |  |  |  |  |
| No | 1715(84.36) |  |  |  |
| Yes | 318(15.64) |  |  |  |
| Follow-up Days | 1604.58 ±1050.16 | | | |

\* P-values are calculated by Fisher's exact test (categorical variables) or Student's t-test (continuous variables).

### Deep-learning based pipeline recognized tissue compartments and extracted whole-slide pathological abnormalities

The tissue compartment detection models were first built on a training set of 11473 image tiles and were applied to 3986 image tiles for accuracy estimation (Table S2). The glomerulus detection model correctly detected 96% of normal glomeruli and differentiated sclerotic glomeruli at True Positive Rate (TPR) of 91% and Positive Predictive Value (PPV) of 93%. The artery detection model detected 94% of arteries and accurately identified 94% of arterial intimal fibrosis regions within lumen. Lastly, the tubule detection model recognized 90% of tubules of differing morphology from frozen sections. Figure S1 demonstrated examples of detected compartments with diverse morphology in frozen tissue. Thus, our deep learning models were capable of detecting kidney tissue compartments as well as abnormalities, despite of freezing artifacts.

Next, we designed a pipeline automatically segmenting a whole slide image into tissue sections, performing compartment detection through pre-trained models to estimate whole-slide abnormalities. Three whole-slide features were extracted in this study: i) Sclerotic Glomeruli% (GS%); ii) Arterial Intimal Fibrosis% (AIF%) per artery was defined by loss% of luminal area by adjusting for the effect of compression or tangential cut. The whole-slide score was determined using the most severely lesioned arteries; iii) Interstitial Fibrosis% (IF%), proportion of the area of enlarged interstitium while minimizing the impact of freezing artifact. Detailed novel strategies of digital feature quantification are descripted in the supplementary methods and Figure S2. Figure 2 shows an example of whole slide prediction, with magnified regions illustrating predicted normality vs. abnormality in the arteries (panel a-c), interstitium (panel d-f) and glomeruli (panel g). Our pipeline successfully avoided freezing artifact affected regions when estimating abnormal interstitium (Panels e. and f.).

**Figure 2.**
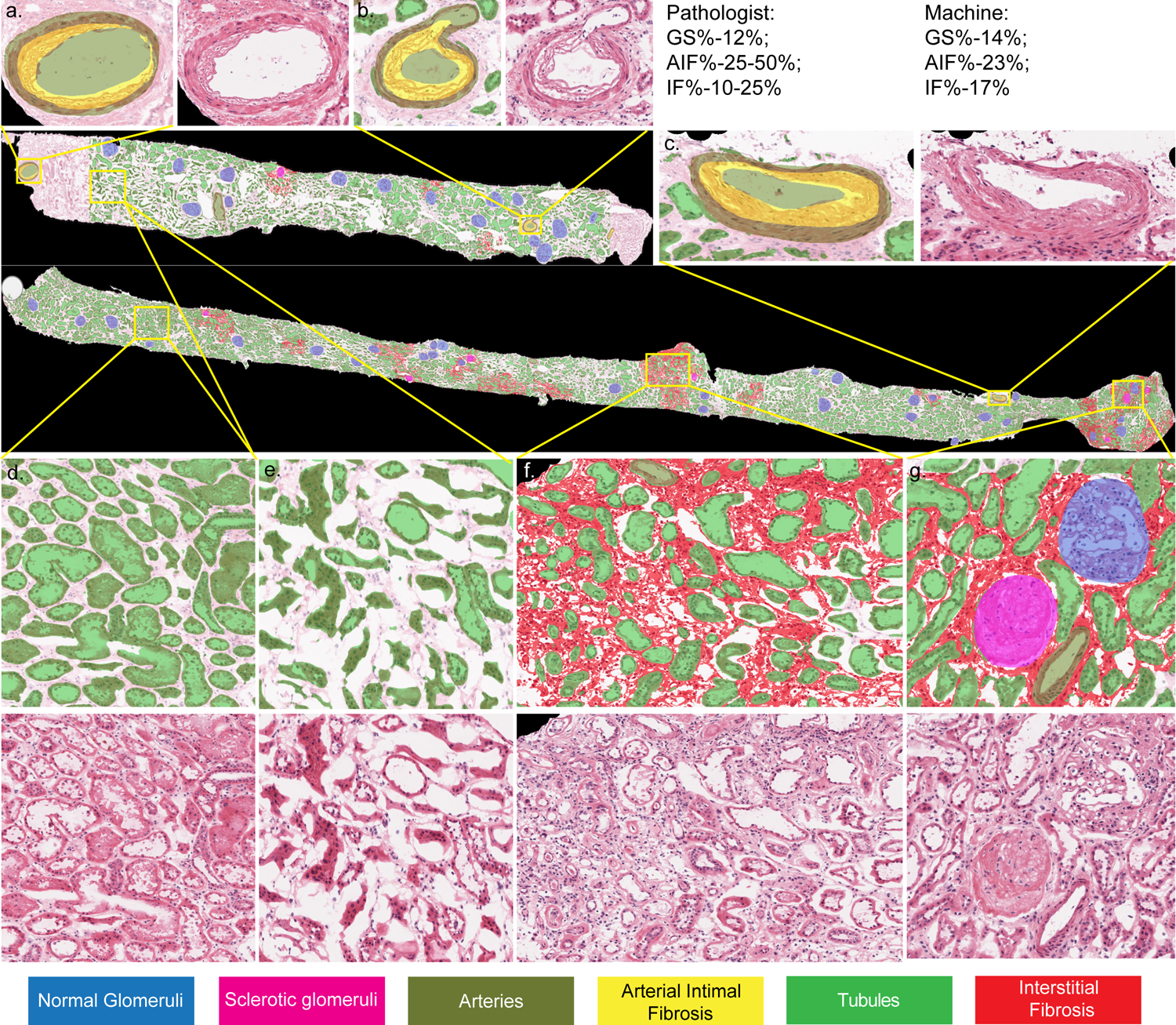
Demonstration of whole slide image prediction. Prediction images of two sections extracted from one whole slide were demonstrated in the middle. We highlighted normal glomeruli in blue, sclerotic glomeruli in magenta, arteries in olive, arterial intimal fibrosis in yellow, tubules in green, and interstitial abnormalities in red. Since the pipeline was designed to perform slide level Interstitial Fibrosis% estimation only in glomeruli enriched region, some regions at left- and right-most side of the top section were excluded during Interstitial Fibrosis% estimation. Sub figures **a-g** demonstrated close-up views of specific regions. **a-c**. Three arteries with arterial intimal fibrosis. **d-e**. Regions with normal tubules with (**e**.) or without (**d**) freezing artifacts. **f**. Region with interstitial fibrosis. **g**. Region with normal and sclerotic glomerulus.

### Digital pathological abnormalities correlated with pathologist scores but were more sensitive to abnormality detection

Our whole slide investigation pipeline was applied to slides from all transplanted and discarded kidneys (n=2431). Compared to the original pathologist reports, we observed significantly strong correlations between our digital scores and pathologist scores (Figure S3). Since the calculation of digital Sclerotic Glomeruli% paralleled pathologist score in a continuous form, it demonstrated the strongest correlation with R=0.75 (Figure S3A). The other two ordinal pathologist scores (Arterial Intimal Fibrosis% and Interstitial Fibrosis%) were also significantly correlated with corresponding continuous digital scores with R=0.43 (*p*=2.3e-109, Figure S3B) and R=0.32 (*p*=5.6e-62, Figure S3C) respectively. Interstitial Fibrosis% had the lowest correlation which could be due to the difficulty of scoring this parameter in frozen sections.

Despite overall agreement between digital and pathologist scores, there were some inconsistencies. We examined transplant outcomes in those inconsistent biopsies there were determined abnormal by digital score only (digitally-abnormal-only), comparing with that in the consistent biopsies scored normal by both methods (normal-by-both). We observed that digitally-abnormal-only group (n=185) demonstrated significantly lower 6m, 12m, 24m estimated glomerular filtration rate (eGFR, mean value: 51.54 vs. 58.35) and higher incidences of graft loss (HR: 2.01 (1.20-3.37) than the consistent normal-by-both group (n=232) (Figure S3D).

These data indicated that our digital scores correlated strongly with pathologist scores but were more sensitive to abnormalities in frozen biopsies, therefore provided more objective and accurate evaluation for those features that were difficult to assess visually, such as AIF% and IF%.

### Digital pathological abnormalities associated with early post-transplant events

To investigate the impact of digital scores in transplant outcomes, we evaluated their associations with post-transplant short-term outcomes (within 24-months) such as eGFR, delayed graft function and acute rejection in 2033 transplanted kidneys. Interestingly, all three digital scores were superior to pathologist scores, and were consistently associated with eGFR at all time points within 24-months in multivariate analyses (Table S3).

Specifically, we observed significant negative correlations of digital scores with 6m, 12m and 24m eGFR for Sclerotic Glomeruli (GS)% (*p*=2.6e-08, *p*=5.8e-11, *p*=2.5e-10) and Arterial Intimal Fibrosis (AIF)% (*p*=1.1e-13, *p*=2.3e-12, *p*=8.1e-10), moderate correlation for Interstitial Fibrosis (IF)% (*p*=2.0e-04, *p*=7.3e-05, *p*=3.4e-04), superior to nearly all corresponding eGFR correlations with pathologist scores (Table S3). Figure S4A shows that the patients who received >10% digital GS% kidneys demonstrated an eGFR decline from 6m to 24m, while the GS%<10% group maintained normal graft function over time. Besides eGFR, digital AIF% was uniquely associated with recipients who required dialysis within the first week post-transplantation (delayed graft function (DGF), *p*=0.003) and from the first week to 12m (slow graft function not defined as DGF, *p*=8.4e-04) (Figure S4B), while AIF% and IF% were also associated with acute rejection at 6m and 12m (Figure S4C-D).

### Digital pathological abnormalities and KDPI were integrated into a composite Kidney Donor Quality Score (KDQS) and were associated with graft loss

We included 1560 kidneys transplanted in 2011-2016 for graft survival analysis, and randomly split in a 1:2 ratio into a n=520 discovery set (D) and a n=1040 validation set (V) while maintaining similar population distribution (Table S4) to derive and validate a composite score for graft survival prediction.

Association of graft loss with individual digital scores was first assessed in the discovery set by cox-regression in multivariate analyses (Table S5). All three digital scores were significantly associated but pathologist scores did not reach statistical significance. Each continuous individual digital score was then converted to an ordinal score (0/1/2, supplementary methods). These ordinal scores were added to derive a combined digital score ranging from 0 to 6 and demonstrated highly significant association with graft loss (*p*=1.9e-05, Table S6).

Besides histopathology, the Kidney Donor Profile Index (KDPI)[36], which incorporates the demographic and clinical characteristics of deceased donors was also associated with graft loss (*p*=0.012, Table S6) in our cohort. Therefore, we converted KDPI to ordinal score (0/1/2) similarly and added to the combined digital score to define a Kidney Donor Quality Score (KDQS, score 0-8, supplementary methods) reflecting the overall quality of the donor kidney. The KDQS was significantly associated with graft loss (*p*=5.6e-06, HR:1.34(1.18-1.52)) in multivariate cox-regression, outperforming all its components (Table S5-7). Such association of KDQS was confirmed in the validation set (*p*=2.8e-05, HR:1.21(1.11-1.33)). Additionally, KDQS was more significantly associated with eGFR (6m: *p*=1.2e-30, 12m: *p*=5.5e-33, 24m: *p*=2.3e-26). When dividing KDQS at the median score value (≥4 and <4), the KDQS≥4 group demonstrated significantly lower survival rate in both the discovery set (Figure 3A, *p*=4.4e-04) and the validation set (Figure 3B, *p*=1.6e-05). Moreover, we found KDQS outperformed Donor Chronic Damage Score[30], Leuven Donor Risk Score[31] and KDPI-alone in association with graft loss in our cohort (Table S7, Figure S5).

**Figure 3.**
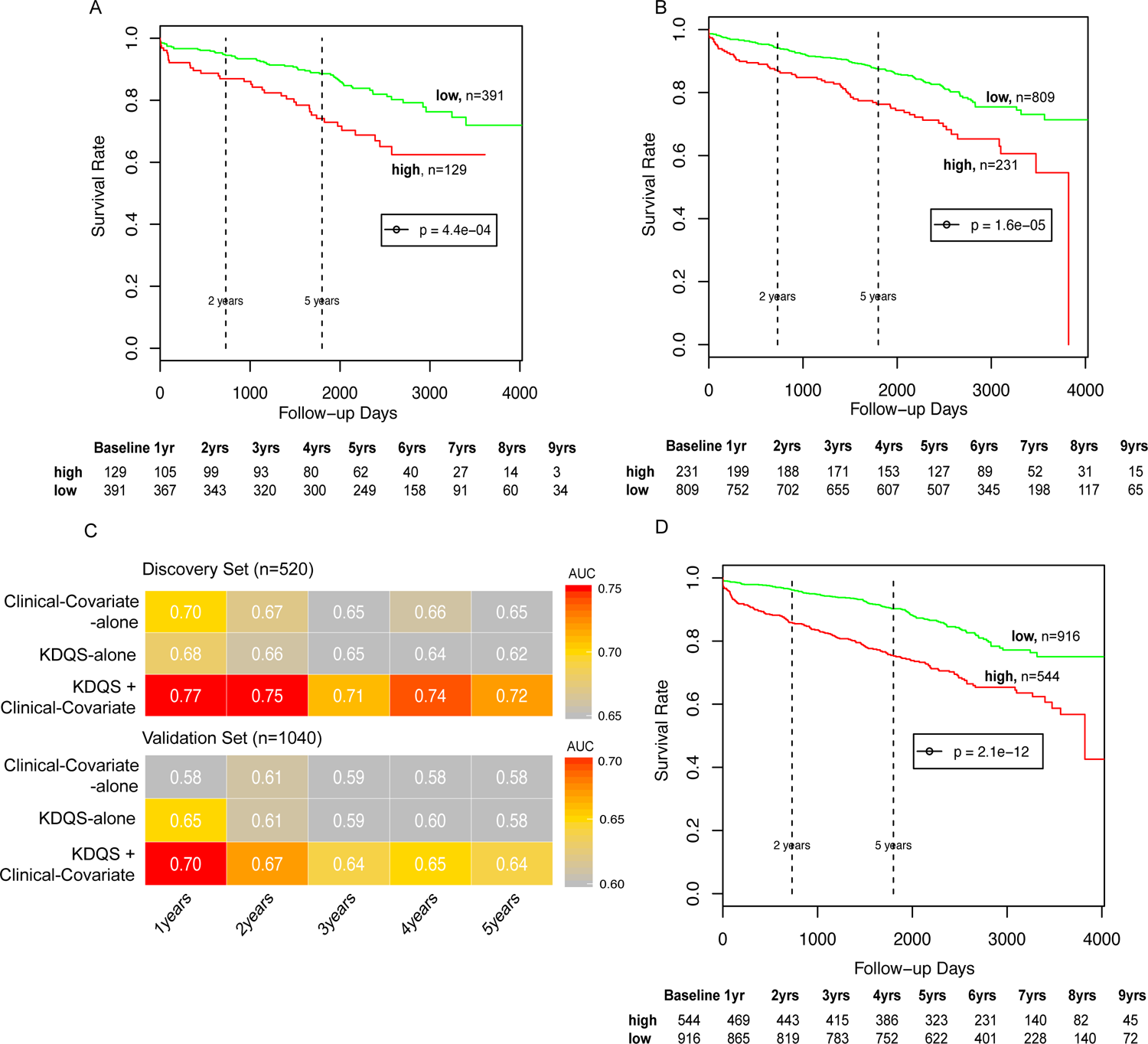
Association of Kidney Donor Quality Score with post-transplant graft loss. **A-B)** Kaplan-Meier curves of graft loss in risk groups stratified by median KDQS value (>4, <4) in the discovery set (**A**) and the validation set (**B**). P-values were calculated by log-rank test. **C)** Heatmaps of the AUCs of Clinical-Covariate-alone, KDQS-alone, and KDSQ + Clinical-Covariate models predicting first 5 years graft loss in the discovery (n=520) and the validation set (n=1040). **D)** Kaplan-Meier curves of graft loss in risk groups stratified by prediction scores from 4-year KDQS + Clinical-Covariate model (cutoff=0.108) in two sets combined. P-values were calculated by log-rank test.

### Kidney Donor Quality Score (KDQS) in combination with peri-transplant and recipient characteristics predicted graft loss

Apart from the donor characteristics, recipient age[37] and race[38], the use of pump perfusion[39, 40]; cold ischemia time (CIT)[41, 42] and induction therapy[43] obtained pre-implantation are also reported to be associated with post-transplant outcomes. We thus first obtained an optimal Clinical-Covariate-alone models by incorporating these known factors to predict graft loss from years 1 to 5 post-transplant in the discovery set (n=520). During stepwise selection process, the use of pump (Y/N), CIT, and induction therapy were selected in first 3 years, while recipient age was only selected from year 4.

We then predicted the graft loss from 1 to 5 years post-transplant by KDQS alone or along with clinical covariates in both discovery (D) and validation (V) sets. The combination of KDQS and clinical factors exhibited a significant prediction improvement, specifically achieved 0.77 and 0.75 in the discovery set and 0.70 and 0.67 in the validation set in 1- and 2-year, respectively (Figure 3C). Since recipient age was identified as a contributor in year 4, we described the ROC curves (Figure S6) and coefficients (Table S8) of 1-year and 4-year graft loss prediction models. The 4-year model predicted intermediate-term graft loss with an overall sensitivity=0.65 and specificity=0.66, and stratified patients into a low-risk group which exhibited stable survival curve with 96% and 90% survival at 2 and 5 years respectively, and a high-risk group which exhibited a progressively reduced graft survival rates from 86% to 75% from 2 to 5 years, further dropping to 65% at 8 years (Figure 3D, *p*=2.1e-12).

### Potential utility of Kidney Donor Quality Score (KDQS) in improving organ utilization

Firstly, to test the hypothesis that KDQS itself could improve organ acceptance rate, we matched 398 kidneys discarded due to biopsy findings (KDQS<4 group: n=110; KDQS≥4 group: n=288) to transplanted kidneys with intermedium-term follow-up (n=1560), and compared the survival curves of discard-matched kidneys to that of the unmatched kidneys (overall remaining transplants). Each discarded kidney was matched (1:1) to its most similar transplanted kidney by minimizing the overall distance based on gender, race, age of donor, KDQS and KDPI (Figure S7), which resulted in 398 discard-matched and 1162 unmatched transplants. We observed no survival difference between the matched kidneys from discarded KDQS<4 group (n=110) and unmatched kidneys (n=1162, *p*=0.871), which suggested that if transplanted, these 110 discarded kidneys could have a similar survival to that of other transplanted kidneys, with 97% and 86% survival at 2 and 5 years respectively (Figure 4A). On the other hand, the matched kidneys from discarded KDQS ≥ 4 group (n=288) showed significantly lower survival rate compared to unmatched kidneys (n=1162, *p*=3.8e-08, Figure 4A).

**Figure 4.**
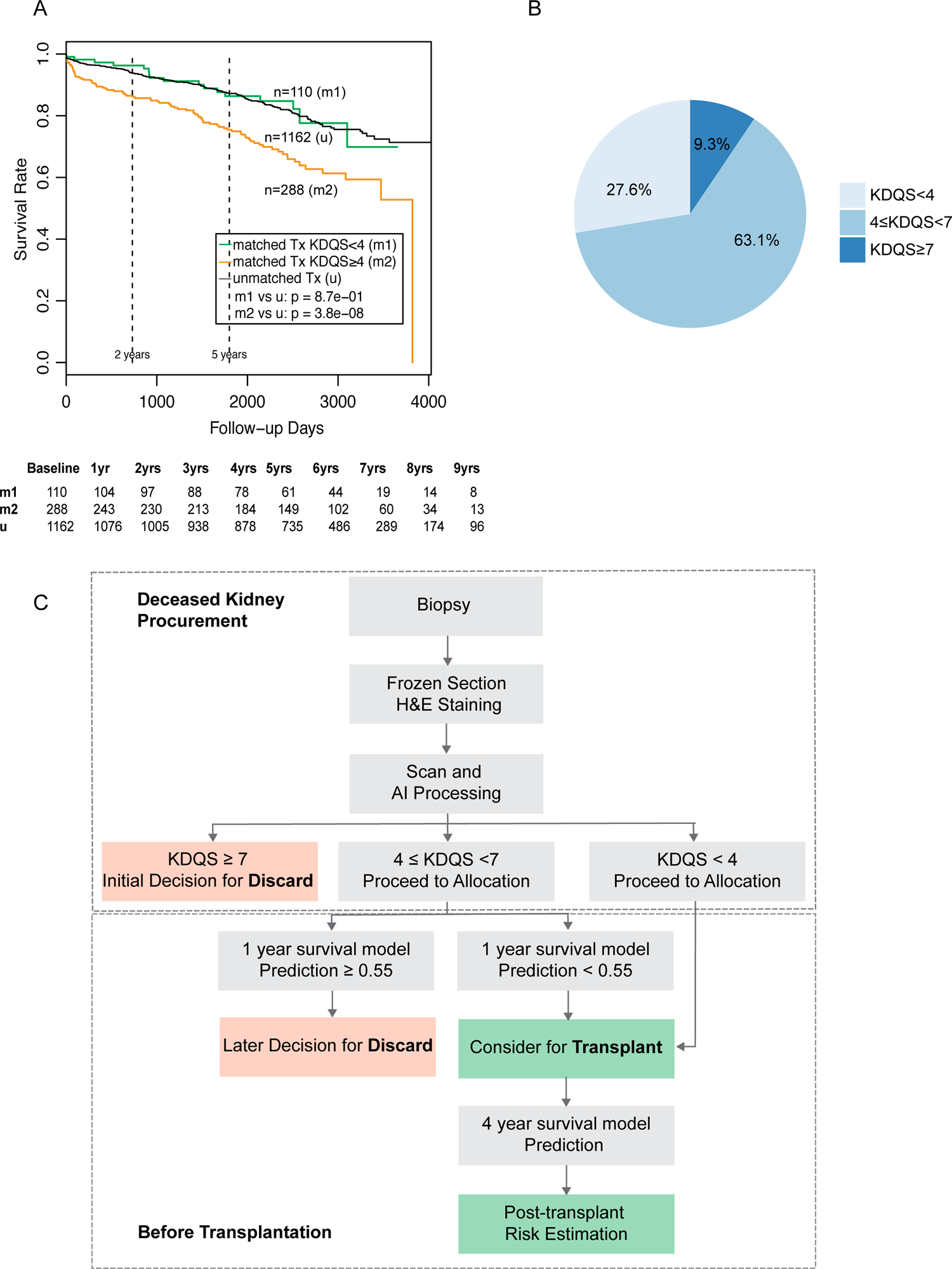
Matching of discarded kidneys due to biopsy findings to transplanted population. **A)** Kaplan-Meier curves of graft loss in matched transplanted kidneys from discarded KDQS>4 group (orange), matched transplanted kidneys from discarded KDQS<4 group (green), unmatched transplanted kidneys (grey) in transplanted intermediate-term follow-up cohort (kidneys transplanted in years 2011-2016). P-values were calculated by log-rank test. **B)** Pie chart demonstrates distribution of KDQS groups in 398 discarded due to biopsy findings kidneys. **C)** Flowchart illustrates the proposed procedure of organ selection based on biopsy findings in clinical practice with the integration of AI and survival prediction models.

Secondly, to determine a possible discard threshold by KDQS or 1-year survival prediction model, we defined a 1-year high risk as severe damage of kidney function (determined by eGFR) or graft loss within 1-year and aimed to obtain a high rate of 1-year high risk above thresholds. We determined KDQS threshold at ≥7 as all components were abnormal with all except one graded in the highest severity, and identified 9 cases from all transplants (0.44%, n=2033), which was equivalent to the proportion determined by severe Donor Chronic Damage Score[30]. Out of these 9 cases, 8 experienced 1-year risks and 1 died within 1-year (Table S9). Accordingly, the threshold of 1-year survival model (Table S8) was observed at the prediction score ≥0.55, which identified additional 4 cases experienced 1-year risks (Table S9). Thus, the two methods together correctly identified 12 out of 13 transplanted kidneys with impaired 1-year graft functions (PPV=0.92) which should have been considered for discard.

Above data showed that application of KDQS could identify those organs discarded but could be potentially transplanted (KDQS<4, 27.6%, Figure 4B), and organs could be discarded in early decision by KDQS alone (KDQS≥7, 9.3%, Figure 4B) or later decision by 1-year survival model, respectively. Both analyses suggested a potential improvement of organ utilization with the advantages of automated, rapid processing and objective scoring.

In clinical practice, the WSI investigation pipeline can be integrated into scanners from various venders and generate digital pathological report either locally or centrally in 30 minutes after biopsy, staining, and scanning procedures. The report and model prediction can be used in conjunction with renal pathologists’ report or independently for organ selection and risk stratification using the proposed procedure described in Figure 4C.

## Discussion

The question of whether to use procurement biopsies has long been debated, as no consistent relationship of pre-transplant biopsy findings with post-transplant graft outcomes has been established[9–15]. On the other hand, “biopsy findings” have become the major reason for deceased door kidney discard[5–7]. Therefore, it is critical to systematically evaluate the utility of procurement biopsies so as to reduce the discard rate and optimize organ risk stratification. We presented here the first large-scale retrospective study involving multiple transplant centers and hospitals of applying artificial intelligence techniques to procurement kidney biopsies to facilitate graft loss risk estimation and organ utilization with superior performance to that of the pathologists.

Our approach has several advantages in evaluating frozen sections. Compared to pre-implantation biopsies, the freezing artifacts in procurement biopsies pose substantial challenges to pathologists during assessment[18–20]. The deep-learning techniques enhanced recognition of tissue compartments in differing morphology in frozen sections. Moreover. instead of subjectively determining the centroid and radius of lumens by drawing tools, our Arterial Intimal Fibrosis% estimation objectively adjusted the effect of tangential cut (Figure S2A) and was unbiasedly evaluated across the slide. Besides, with ice crystals and inflammation approximation, our Interstitial Fibrosis% estimation was able to eliminate most of regions affected by freezing artifacts which appeared to be abnormal (supplementary methods). Those measures ensured the unbiased evaluation and could led to improved association with graft outcomes.

In contrast to previous studies that only glomerulosclerosis was associated with graft outcomes in procurement biopsies[44], all three digital features (GS%, AIF%, IF%) defined in our study were strongly associated with post-transplant eGFR and graft loss, and digital AIF% was also associated with various other outcomes at high significance. Perez-Gutierrez et al. recently reported AIF% in reperfusion but not procurement biopsy correlated with GFR[45]. Two reasons raised by author were: 1) Unlike reperfusion biopsies, their procurement biopsies were evaluated by on-call pathologists. 2) the preference of wedge biopsies for procurement biopsies, which was less suited for vasculature evaluation[45]. Others also found 1) only evaluation by renal pathologists correlated with graft outcomes and evaluation by on-call pathologists tended to lead to more discards, 2) needle biopsies perform much better in the evaluation of vascular lesions[19]. Given our large sample size, the use of needle biopsies, and deep-learning models trained under guidance of experienced renal pathologists, we believe our pipeline overcame these issues and improved the prognostic value of procurement biopsies in extracting lesion features (particularly AIF%).

Furthermore, by incorporating three digital scores with KDPI, we derived a Kidney Donor Quality Score which outperformed all components and public scores in associating with graft loss. Because KDPI included most donor characteristics contributing to discard[6], KDQS theoretically represented overall quality of donor kidney. KDQS itself can be used for initial screening after organ recovery: 1) poor quality kidneys (KDQS≥7) may be discarded and low KDQS (<4) kidneys may be more suitable for transplant. 2) when a certain level of survival risk was otherwise acceptable, intermediate KDQS (4≤KDQS<7) kidneys may also be accepted. Then, before transplantation, KDQS + Clinical-Covariate models can assist in last-minute selection (1-year model) and intermediate-term risk evaluation to identify patients needing monitoring or preventative measures (4-year model). Of a note, it is known that deceased donor kidneys in US tend to have significantly better conditions than that in other countries[4], thus the number of identified discards is expected to be low in our transplanted cohort.

Our study has some limitations. Firstly, the performance of survival prediction models remains to be improved. Although the original paper [31] reported the Leuven Donor Risk Score predicted survival with AUC of 0.7-0.8, Hall et al.[46] reported a lower AUC of 0.61 in a larger cohort. Also, all public scores were derived on pre-implantation biopsies. When evaluating in our cohort, both public scores underperformed KDQS in graft loss association. Moreover, we noticed that the estimation of IF% remained less accurate than other two digital scores, resulting in weaker association with outcomes. More training of tubular changes in in frozen sections will be needed to improve the IF% estimation, which could further improve performance of survival prediction models. Secondly, although we demonstrated our approach can predict post-transplant graft loss based on large-scale investigation from multiple transplant centers and hospitals nationwide, the slides were obtained in 2011-2015. Thus, we are collaborating with transplant centers and hospitals patterned with the Organ Procurement and Transplantation Network (OPTN) to prospectively investigate the influence of our approach in current transplant or discard procedures. Thirdly, the pipeline itself is still command-line based. We are working to develop user-friendly interface and integrate it into various types of scanners for more reliable and cost-efficient pathological assessment of procurement biopsies, especially for those transplant centers lacking experienced renal pathologists.

In conclusion, we constructed deep-learning based models detecting tissue compartments in H&E stained slides of frozen procurement biopsies form deceased donor kidneys, and further developed a pipeline extracting pre-transplant pathological abnormalities. We also derived a composite KDQS and survival prediction models which could facilitate risk stratification and organ utilization, and potentially reduce unnecessary organ discard.

## Author Contributions

Z.Y. designed and performed computational analyses, and drafted the paper. C.X. annotated slides and managed clinical and image data. M.C.M contributed to analysis design and paper editing. F.T., B.V. were involved in pathological/clinical data analysis and interpretation; A.S., M.G, and R.B scanned the slides; Z.S., C.W., M.C. and W.W. were involved in slide and image data management; K.L. and J.Z. were involved in programming for image processing and object detection; A.K. were involved in clinical data analysis; P.C., L.G., P.O.C, M.N., R.S. and R.B.C were involved data interpretation and edited the paper. S.W. provided pathological slides and report, supervised pathology annotation and interpretation and edited the paper; F. S. conceptualized the study, supervised pathology annotation and interpretation; W. Z. conceptualized and designed this study and edited the paper. All authors reviewed and approved the manuscript.

## Data Sharing

Individual patient data is not allowed to make publicly available based on Data Usage Agreement (DUA) between OPTN and investigators. The code may be made available to qualifying researchers by requesting to corresponding authors. Proposals will be reviewed by the investigators and collaborators based on scientific merit. If the proposal is approved, the data will be shared through a secure data transfer site.

## Disclaim regarding OPTN data

The data reported here have been supplied by UNOS as the contractor for the OPTN. The interpretation and reporting of these data are the responsibility of the author(s) and in no way should be seen as an official policy of or interpretation by the OPTN or the U.S. Government.

## Acknowledgement

We thank Scientific Computing Division at the Icahn School of Medicine at Mount Sinai for providing computational resource. We thank Pathology Core at Mount Sinai for slides scanning. This work was supported by Biocomputation Fund (02435913) from Department of Medicine and i3 Genesis award from Mount Sinai Innovation Partner (MSIP) of Icahn School of Medicine at Mount Sinai (AGR-25489). We thank Miss. Denise Tripp and Miss. Aparna Sadavarte from UNOS organization for coordinating the access to UNOS database.

