## Supplementary files for "Deep-learning Based Pathological Assessment of Frozen Procurement Kidney Biopsies Predicts Graft Loss and Guides Organ Utilization: A Large-scale Retrospective Study"

### Supplementary Appendix

### Supplementary Methods

#### Image annotation and pre-processing

We annotated multiple sections from 100 H&E stained slides for tissue compartment prediction using ASAP software (https://computationalpathologygroup.github.io/ASAP/) under the guidance of experienced renal pathologists. Each section was outlined by a boundary and then glomeruli, arteries and tubules within the boundary were further annotated. Sclerotic glomeruli and the layers of arterial intimal fibrosis were specifically annotated as separate abnormal groups. The raw sections were divided into 15459 fixed-sized image tiles at 10x objective-power and then transformed by data augmentation process[1] including position shifting, rotating, flipping, perspective transforming, color transferring, contrast transforming, or noise feeding. Every fixed-sized tile image was paired with ground-truth classification information for model evaluation. These two types of images were served as input in model construction process.

#### Deep learning model construction

We split annotated images roughly in the ratio 2:1 into training set (n=11473 images) and testing set (n=3986 images). Training set were used for model construction and testing set were used for final evaluation and were kept untouched during training process. We performed 5-fold cross-validation within training set thus divided the training set into 5 equal sized portions. During each model training process, we used 4/5 of samples as training set and the left-out 1/5 of samples as validation set to tune the model. As a result, 5 separate base models were created based on 5 partitions and the final prediction were made by aggregating results from each base model.

We constructed compartment detection models for glomeruli (including sclerotic glomeruli), arteries (including the layers of arterial intimal fibrosis) and tubules separately using Mask R-CNN structure[2, 3]. Specifically, the glomerulus detection model was trained for 120 epochs with a batch size of 10 using 1024x1024 pixels input images and differentiated normal vs. sclerotic glomeruli; the artery detection model was trained for 180 epochs with a batch size of 10 using 1024x1024 pixels input images and detected all arteries as well as thickening layers in the lumen if available; the tubule detection model was trained for 150 epochs with batch size of 10 using 1024x1024 pixels input images. The detection model used RestNet-101 as backbone and was tuned based on pre-trained weights from MS COCO dataset[4]. We applied GSD optimizer with learning rate of 0.001 and momentum of 0.9. Total loss was calculated as the sum of loss of RPN classifier, RPN bounding box, MRCNN classifier, MRCNN bounding box, and MRCNN mask, where cross-entropy was used for classification problems and smooth L1-loss was used for bounding box refinement. Additionally, in order to more accurately estimate interstitial area, we also constructed a segmentation model of tubule prediction using U-net structure[5, 6]. The segmentation model was constructed using 3 down-sampling layers and 3 up-sampling layers and the first layer contained 32 feature maps. ADAM optimizer was chosen for weight updating at learning rate of 0.001 and cross-entropy was used for loss function. Best epoch/model was determined by evaluating loss from each training and validation partition after 100 epochs. We obtained full prediction of tubules by overlaying results from both detection and segmentation model. By subtracting area of glomeruli, arteries and tubules, the remaining area was therefore determined as interstitial area.

Accuracies were measured by True Positive Rate (TPR) and Positive Predictive Value (PPV) and general $F_\beta$ score[7] where $\beta$=2.

The GPU machine we used was equipped with 36 Intel(R) Xeon(R) W-2195 CPUs (18 cores), 128 GB Memory, and 4 GPUs of Quadro RTX 8000. All processes ran on Ubuntu 18.04 system.

**Automatic extraction of tissue section**

The digital color image of whole slide was first converted to grayscale image. These grayscale values were then inverted by complement filter so that background (originally white) were inverted to black color. Binary image was later generated by application of Otsu thresholding[8]. Closing morphology operator[9] was performed to remove small background noises. The boundary of whole slide was therefore extracted by retrieving top-level base contours[10]. One base contour represented one tissue section and most slides contained >1 contours/sections. All pixels outside of base contours were regarded as background therefore were defaulted to black color.

**Estimation of lesion scores**

**Percentage of sclerotic glomeruli**

Percentage of sclerotic glomeruli was calculated as the number of sclerotic glomeruli divided by total number of glomeruli.

**Percentage of arterial intimal fibrosis**

In general, the percentage of arterial intimal fibrosis per artery was defined by the loss of luminal area in its original circular structure and this lesion score per slide was determined in the most severely affected artery [11]. However, in practice, most of the lumens are compressed, some of the remaining lumens are deviated from center of original circular structure or incomplete due to the tangential cut. Moreover, the size of arteries/lumens varies across the slide and over-estimation of intimal fibrosis are more likely to occur in small arteries/lumens.

We developed a novel approach to overcome above issues. As illustrated in Figure S2A, we first reverted each compressed lumen to its original circular form based on the perimeter of original luminal area. The centroid and radius of original lumen ($r_0$) was thus determined (red circle). Then the centroid and radius ($r_1$) of remaining luminal area were also determined by exclusion of detected intimal fibrosis region from original lumen (white area). Here the remaining luminal area indicates narrowed lumen as a result of arterial intimal fibrosis, while the original luminal area was regarded as the convex hull[12] of intimal fibrosis region (yellow area) and non-fibrotic remaining lumen (white area). Next, the shift of centroid ($d$) was obtained by calculating distance between two centroids and then added to the radius of remaining lumen ($r_1 + d$) to determine the radius of adjusted remaining lumen in consideration of tangential cut (Figure S2A right panel). Finally, the area of original and adjusted remaining lumen was calculated using formula: $A = \pi r^2$ and arterial intimal fibrosis percentage was determined as: (Area of original lumen – Area of adjusted remaining lumen)/Area of original lumen ($[\pi r_0^2 - \pi(r_1 + d)^2]/\pi r_0^2$). This process was repeated for every detected artery and a list of arterial intimal fibrosis percentage was thus obtained for each slide.

In order to fairly estimate this lesion score over whole slide, we followed four steps: Firstly, we filtered out extremely small arteries which may cause biased estimation. Secondly, all remaining arteries were ranked according to their intimal fibrosis percentage values. Top three arteries were selected as determining arteries for whole slide estimation if available, otherwise all arteries with intimal fibrosis percentage >0 were selected. Thirdly, we calculated arithmetic mean and weighted mean of selected intimal fibrosis percentage respectively where the later one was weighted by the size of selected arteries. The final slide-wide arterial intimal fibrosis percentage was determined by averaging previous two means to assign more influence to relatively large arteries and meanwhile avoid biased estimation in huge arteries. On a side note, in the case of incomplete artery or artery with unclosed intimal fibrosis hollow, we penalized the calculated percentage by multiplying a factor of 0.6 to lower its chance to be selected as determining artery.

**Percentage of interstitial fibrosis**

Interstitial area was only examined in glomeruli enriched regions (cortex sections) with the aid of glomeruli detection to avoid over-estimation in medulla sections. To estimate slide-wide percentage of interstitial fibrosis, we first introduce the concept of Region of Interest (ROI) window to identify local abnormal regions with respect to interstitium. Given a whole slide prediction image, we applied a 256x256 pixels unit window sliding over the image with stride of 64 pixels. Within each unit window, we examined following metrics: percentage of tubules, percentage of normal glomeruli, percentage of sclerotic glomeruli, percentage of arteries. The unit window was defined as Region of Interest if it contained sclerotic glomeruli (>0% sclerotic glomeruli prediction), or it was not close to large arteries (no large artery prediction in surrounding area) and was not predominantly occupied by normal glomeruli (<30% normal glomeruli prediction) and tubules (<40% tubules prediction).

The artifacts (particularly tubular shrinkage, ice crystal formation) resulting from freezing procedure appeared frequently over slide and could lead to pathological misinterpretations and misdiagnose especially regarding estimation of interstitial area[13, 14]. Thus, we evaluated two additional metrics: percentage of ice crystals and count of nuclei in interstitium, to eliminate possible false positive Region of Interest windows. In frozen tissues, tubular cells are usually compressed by expanding ice crystals with appearance similar to bubbles in background color. Therefore, the extracted white background pixels surrounding tubules were added to prediction of tubules. Remaining white background pixels were summed and divided by sum of interstitial pixels (pixels in tissue but not predicted as any object or artifacts) and white background pixels within interstitium to obtain a percentage of ice crystals per unit window. Moreover, since renal inflammation and fibrosis often occurs together[15], approximation of the number of inflammatory cells also assist with artifact elimination. The cell nuclei were stained and distinguished from other structures in purplish blue color in H&E slides[16], we extracted and counted contours[10] of nuclei in interstitial area (or precisely, not in other detected compartments such as glomeruli, tubules, arteries, or ice crystals). This number was used to approximate inflammation within interstitium within the unit window. The previously identified Region of Interest windows were re-examined with updated percentage of tubules and artifact related metrics. Final Region of Interest window also contained < 30% ice crystals within window or <40% ice crystals within interstitium, and > 50 nuclei within interstitium.

We scanned all glomeruli enriched regions (cortex sections) for each slide using the unit window and summarized interstitial pixels within Region of Interest (ROI) windows, then divided by interstitial pixels in whole slide to obtain slide-wide percentage of interstitial fibrosis.

**Derivation of Kidney Donor Quality Score**

Three digital pathological scores and KDPI were included for composite score calculation. Firstly, by following a similar concept as previously-published strategy in searching for an optimal cutoff of glomerulosclerosis when generating Donor Chronic Damage Score [17], two optimal cutoffs were searched against lower (≤median value) and upper range (>median value) of each digital score and KDPI through logistic regression of graft loss prediction from 1 to 7 years. We determined the upper-tertile as the follows: i) A search of sub-optimal threshold was first performed within upper range of the score with the most significant association through logistic regression of 1-year graft loss prediction. ii) Repeat previous step for 2- and 3-year graft loss prediction. iii) The optimal threshold was then determined as the maximum value of list of identified sub-optimal thresholds of the score. Similarly, we determined the lower-tertile as the follows: i) A search of sub-optimal threshold was first performed within lower range of the score with the most significant association through logistic regression of 4-year graft loss prediction. ii) Repeat previous step for 5-, 6- and 7-year graft loss prediction. iii) The optimal threshold was then determined as the minimum value of list of identified sub-optimal thresholds of the score. Each component score was then converted into integer number ranging from 0 to 2 based on tertile cutoffs, and final composite Kidney Donor Quality Score was determined as the sum of four component scores as shown in formula (1).

$$KDQS = gs + aif + if + kdpi, \ \ where \qquad (1)$$

$$gs = \begin{cases} 0 & GS\% \leq 0.03 \\ 1 & 0.03 < GS\% \leq 0.09 \\ 2 & GS\% > 0.09 \end{cases}; \qquad aif = \begin{cases} 0 & AIF\% \leq 0.11 \\ 1 & 0.11 < AIF\% \leq 0.39 \\ 2 & AIF\% > 0.39 \end{cases}$$

$$if = \begin{cases} 0 & IF\% \leq 0.1 \\ 1 & 0.1 < IF\% \leq 0.39 \\ 2 & IF\% > 0.39 \end{cases}; \qquad kdpi = \begin{cases} 0 & KDPI \leq 0.5 \\ 1 & 0.5 < KDPI \leq 0.76 \\ 2 & KDPI > 0.76 \end{cases}$$

**Association of Graft Outcomes and Prediction of Graft Loss**

When evaluating association of donor biopsy scores with graft outcomes or graft loss, multivariate linear regressions or cox-regressions were performed respectively by adjusting for recipient related factors: gender, race, Estimated Post Transplant Survival (EPTS) score (combination of recipient age, prior transplant and years on dialysis)[18], and peri-transplant characteristics which are reported to be associated with post-transplant outcomes: use of pump perfusion[19, 20]; cold ischemia time (CIT)[21, 22] and induction therapy[23].

Prediction of graft loss at time point t were performed by logistic regression. Let $T$ denotes follow-up days of a patient, at time $t$; a case was defined as the patient having lost the graft at $T \leq t$; a control was defined as the graft having survived through $t$ ($T>t$). Stepwise selection of recipient demographic and peri-transplant characteristics using Akaike Information Criterion (AIC) [24] was first carried out to determine the Clinical-Covariate-alone model at each year. Digital score was then added to construct the combined models. Sensitivities and specificities were calculated based on cutoffs determined by shortest distance from ROC curves to point (0,1) in the discovery set.

**The Matching of Discarded Kidneys**

The matching of discarded kidneys to transplanted kidneys based on KDQS and demographic features (donor gender, age, race and KDPI) was performed by the R package “MatchIt”[25]. A 1:1 nearest neighbor matching was used thus each discarded kidney was paired with a transplanted kidney that had the closest propensity score to it. The propensity score was estimated based on donor characteristics in the logistic regression model.

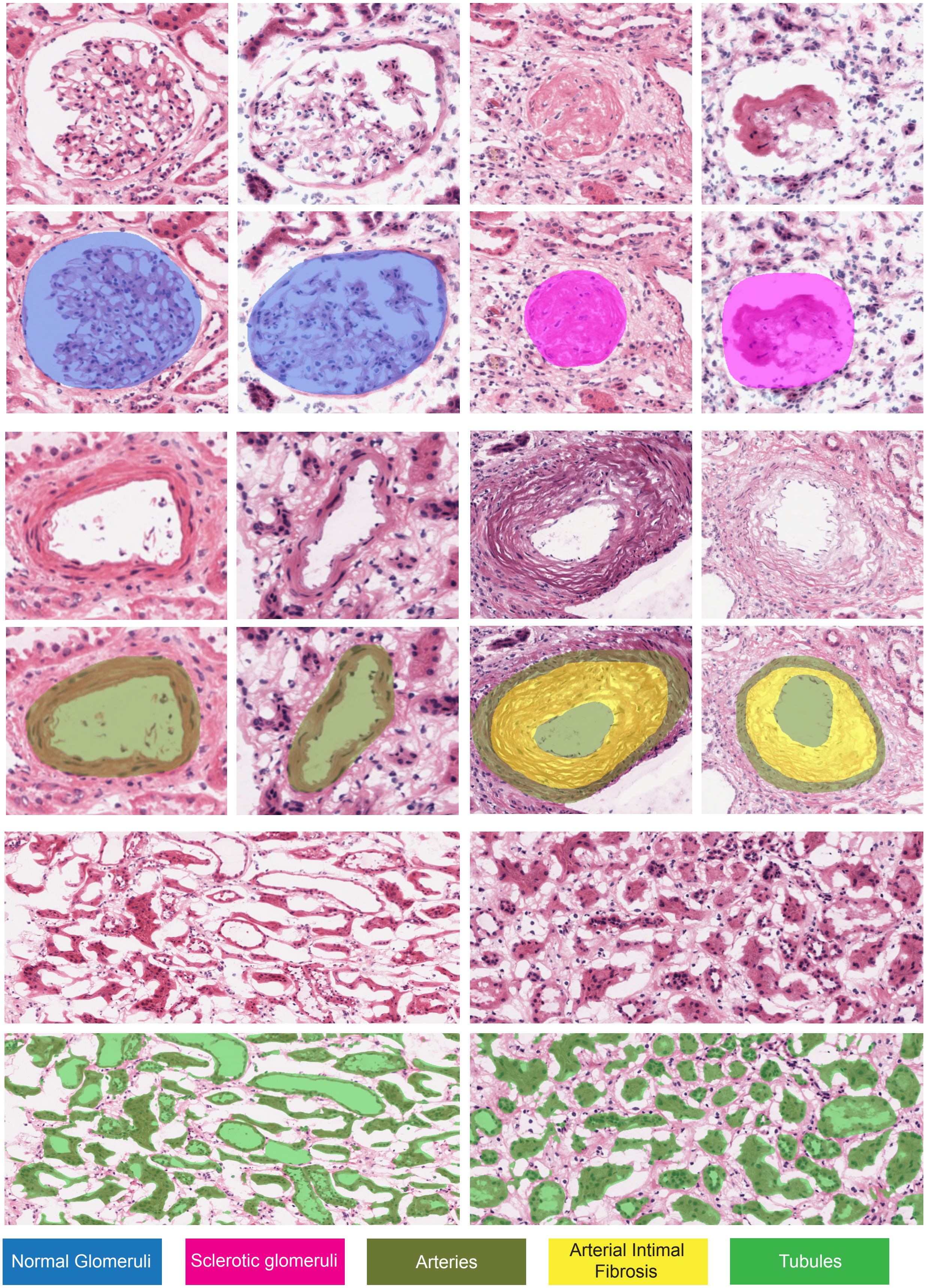

**Figure S1. Examples of kidney tissue compartment detection.** Demonstration of original (upper) and predicted (lower) kidney tissue compartments from deep-learning detection models. We higlighted normal glomeruli in blue, sclerotic glomeruli in magenta, arteries in olive, arterial intimal fibrosis in yellow, tubules in green.

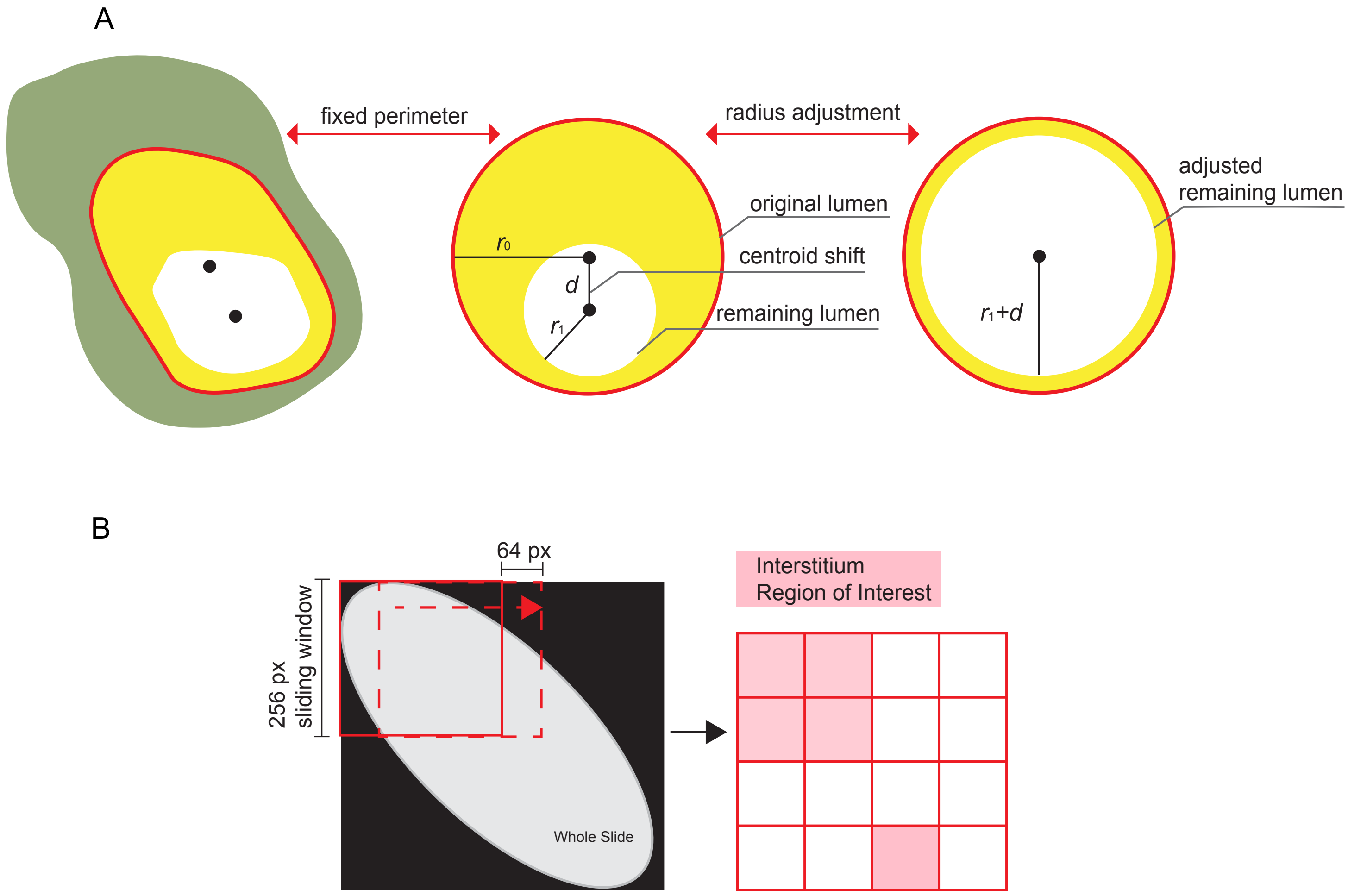

**Figure S2. Illustration of digital Arterial Intimal Fibrosis% and Interstitial Fibrosis% estimations. A)** Illustration of Arterial Intimal Fibrosis% calculation per artery. We reverted each detected lumen to its circular form and obtained the centroid and radius ($r_0$) of original lumen. Then the centroid and radius ($r_1$) of remaining luminal area were also determined by exclusion of detected intimal fibrosis region from original lumen (white area). Next, the shift of centroid ($d$) was obtained by calculating distance between two centroids and the radius of adjusted remaining lumen was calculated as $r_1+d$. Finally, the Arterial Intimal Fibrosis% per artery was calculated as $[(\pi r_0^2)-\pi(r_1+d)^2]/\pi r_0^2$. The right most panel indicates final boundary of remaining lumen after adjusting for the effect of tangential cut. **B)** Illustration of slide level Interstitial Fibrosis% calculation. We applied a fixed sized unit window sliding across the slide and evaluated interstitial fibrosis related metrics within each window. The unit window was defined as Region of Interest (ROI) if it contained sclerotic glomeruli, or it was not close to large arteries and was not predominantly occupied by normal glomeruli and tubules. Final Region of Interest (ROI) windows were also re-examined to minimizing effect of freezing artifacts. These ROI windows were then assembled into global interstitial abnormality map for slide level Interstitial Fibrosis% calculation.

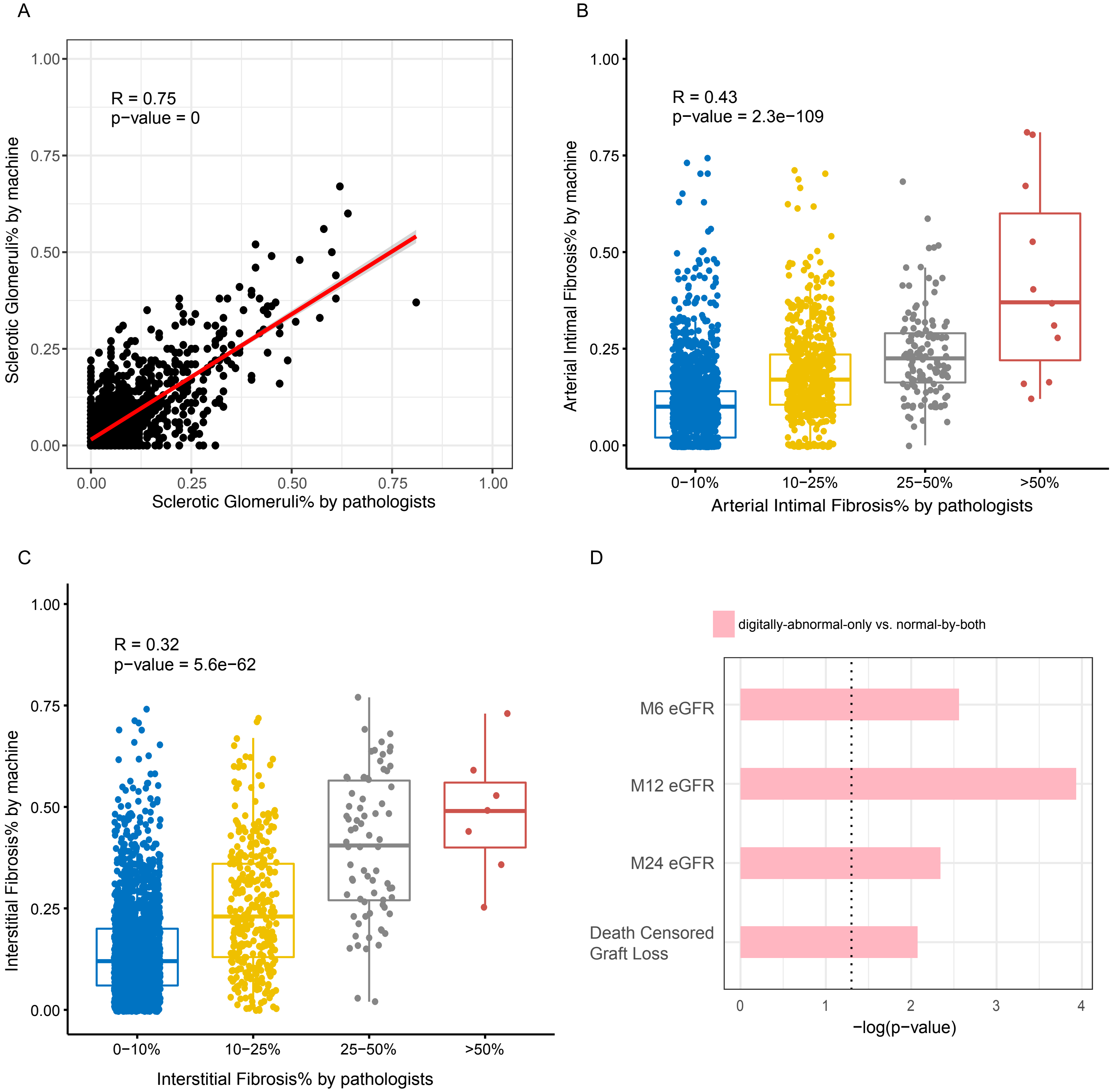

**Figure S3. Correlation of digital pathological scores with the pathologist scores.** The scatter plot of digital vs. pathologist scores for **A**) Sclerotic Glomeruli%. The box plots of digital vs. pathologist scores for **B**) Arterial Intimal Fibrosis% and **C**) Interstitial Fibrosis%. **D**) The bar chart shows –log(p-value) of comparisons of graft outcomes in digitally-abnormal-only group vs. consistent normal-by-both group after adjusting for recipient and peri-transplant information in multivariate analyses. The associations with eGFR were assessed by linear regression, the associations with graft loss were assessed by cox regression. Bars surpassed the dotted line indicate significant associations (p-value<0.05).

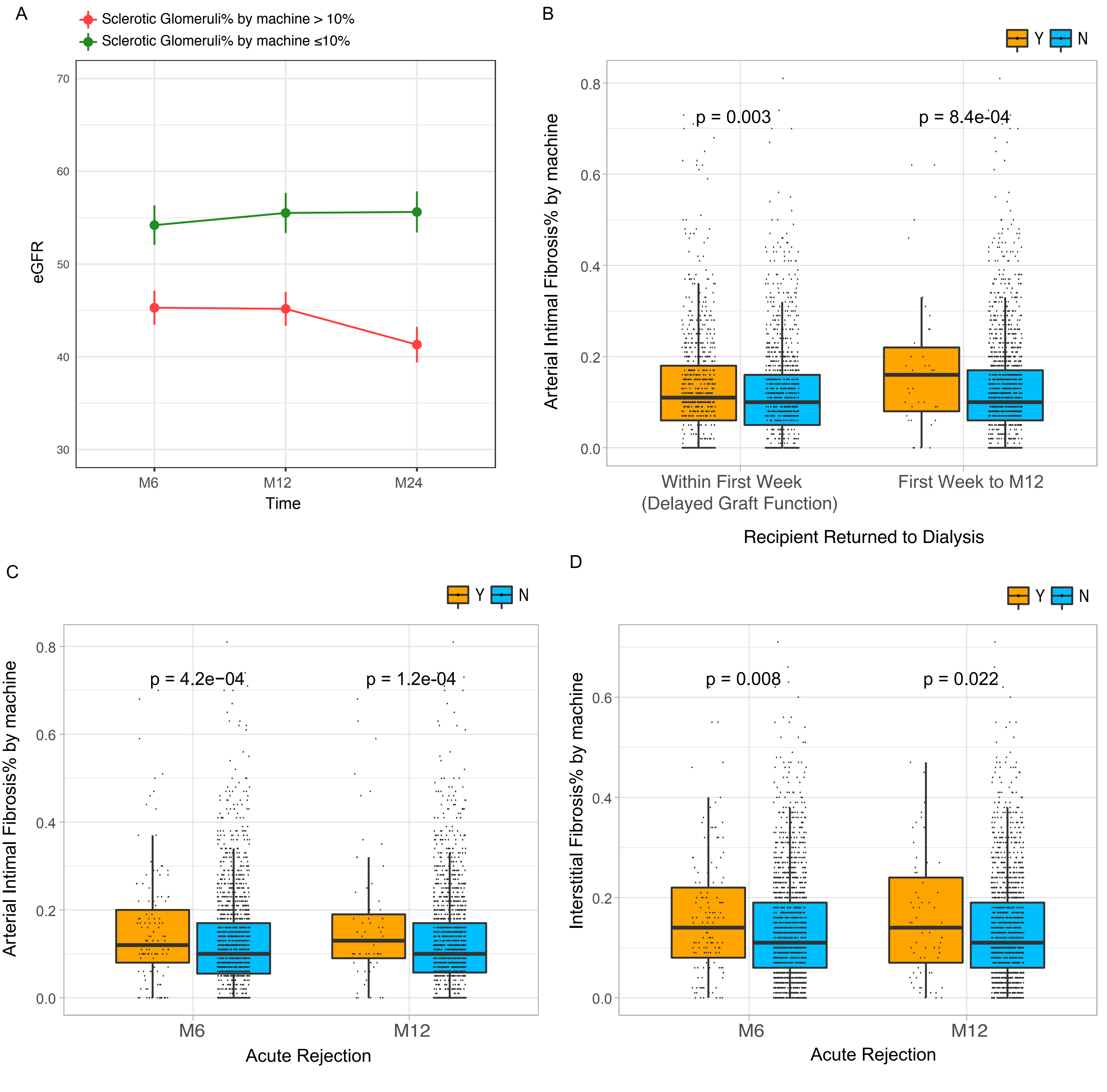

**Figure S4. Association of digital abnormality scores with post-transplant early graft outcomes. A**) Average eGFR values over time within 24m post-transplant for digital Sclerotic Glomeruli%>10% and ≤10% group. Error bars represent 0.1x standard deviation from mean values. **B**) Association of digital Arterial Intimal Fibrosis% and recipient returned to dialysis within first week (delayed graft function) and first week to 12m post-transplant. P-values are from linear regression by adjusting for recipient and clinical information. **C-D**) Association of digital Arterial Intimal Fibrosis% (**C**) and digital Interstitial Fibrosis% (**D**) with acute rejection at 6m and 12m. P-values are from linear regression by adjusting for recipient and clinical information.

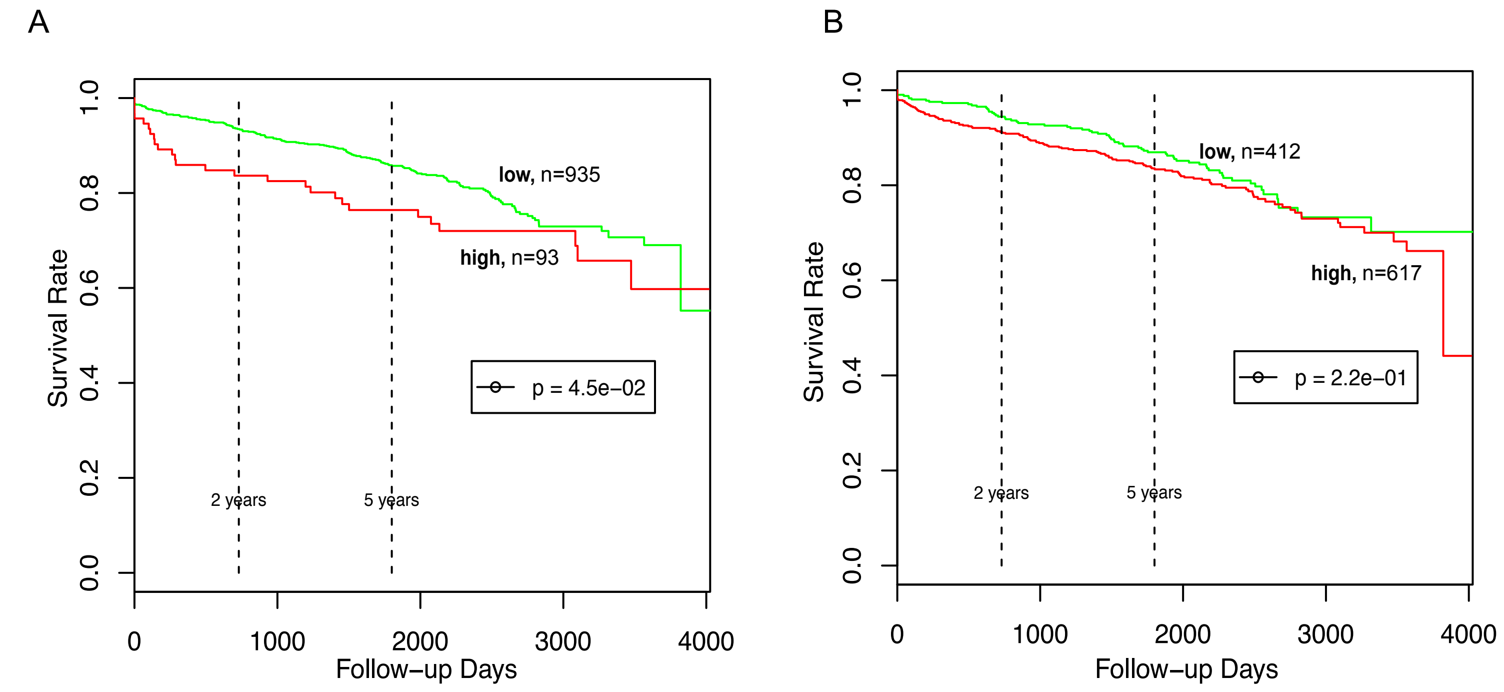

| | Baseline | 1yr | 2yrs | 3yrs | 4yrs | 5yrs | 6yrs | 7yrs | 8yrs | 9yrs |
|---|---|---|---|---|---|---|---|---|---|---|
| high | 93 | 77 | 74 | 70 | 63 | 57 | 46 | 30 | 25 | 13 |
| low | 935 | 863 | 806 | 746 | 687 | 568 | 383 | 217 | 123 | 67 |

| | Baseline | 1yr | 2yrs | 3yrs | 4yrs | 5yrs | 6yrs | 7yrs | 8yrs | 9yrs |
|---|---|---|---|---|---|---|---|---|---|---|
| high | 617 | 555 | 524 | 479 | 440 | 366 | 257 | 150 | 93 | 51 |
| low | 412 | 385 | 356 | 337 | 310 | 259 | 172 | 97 | 55 | 29 |

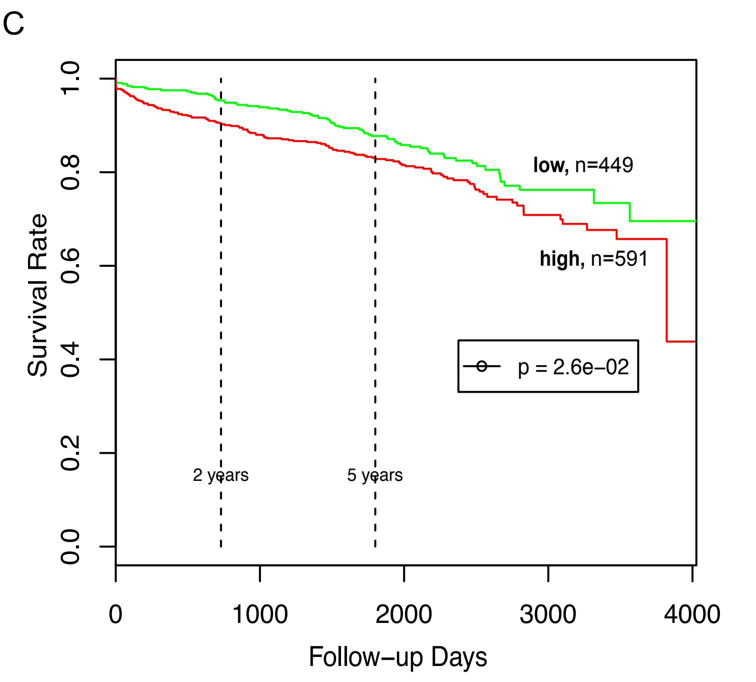

| | Baseline | 1yr | 2yrs | 3yrs | 4yrs | 5yrs | 6yrs | 7yrs | 8yrs | 9yrs |
|---|---|---|---|---|---|---|---|---|---|---|
| high | 591 | 530 | 496 | 451 | 417 | 343 | 244 | 139 | 83 | 45 |
| low | 449 | 421 | 394 | 375 | 343 | 291 | 190 | 111 | 65 | 35 |

**Figure S5. Association of public composite scores with post-transplant graft loss.** Kaplan-Meier curves of graft loss in risk groups stratified by public composite scores: Donor Chronic Damage Score (**A**), Leuven Donor Risk Score (**B**), KDPI (**C**). P-values are calculated by log-rank test. Thresholds of scores was determined at median score value (Donor Chronic Damage Score: 2, Leuven Donor Risk Score: 65, KDPI: 0.5).

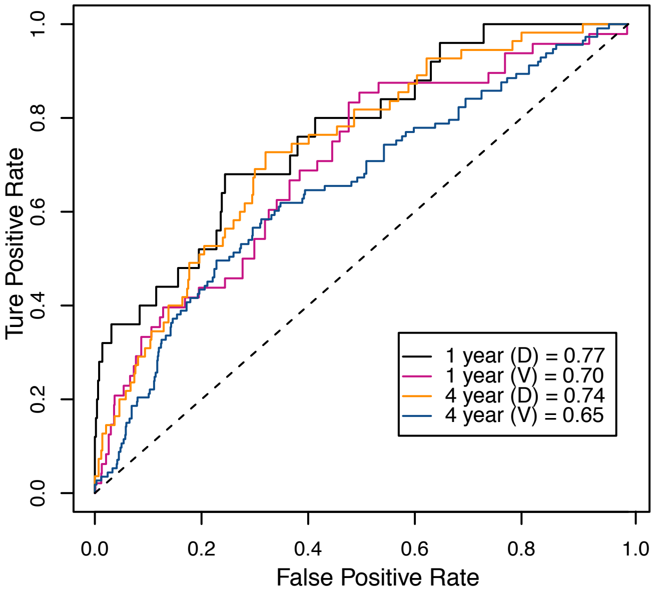

**Figure S6. ROC curves of 1- and 4-year KDQS + Clinical-Covariate model predicting graft loss in discovery and validation set.**

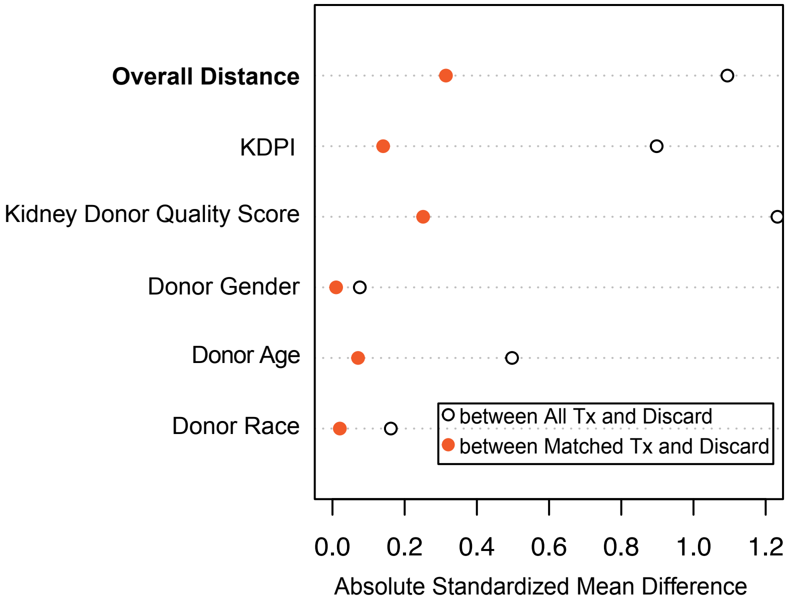

**Figure S7. Matching of discarded kidneys due to biopsy findings to transplanted population.** The plot summarizes covariate balance between discarded and transplanted population before (hollow circle) and after (solid circle) matching. The first "Overall Distance" variable represents overall balance.

**Table S1.** Distribution of Transplant Centers where kidneys were transplanted.

| Transplant Center | Number of Kidneys transplanted |
|---|---|
| ALUA-TX1 University of Alabama Hospital | 2 |
| ARUA-TX1 UAMS Medical Center | 3 |
| AZMC-TX1 Mayo Clinic Hospital Arizona | 7 |
| AZUA-TX1 Banner University Medical Center-Tucson | 2 |
| CACS-TX1 Cedars-Sinai Medical Center | 3 |
| CAGH-TX1 Scripps Green Hospital | 1 |
| CAIM-TX1 University of California Irvine Medical C | 1 |
| CALL-TX1 Loma Linda University Medical Center | 2 |
| CAPM-TX1 California Pacific Medical Center-Van Nes | 1 |
| CASD-TX1 University of California San Diego Medica | 2 |
| CASF-TX1 University of California San Francisco Me | 4 |
| CASH-TX1 Sharp Memorial Hospital | 1 |
| CASM-TX1 University of California Davis Medical Ce | 7 |
| CASV-TX1 St. Vincent Medical Center | 1 |
| CAUC-TX1 University of California at Los Angeles M | 1 |
| CAUH-TX1 Keck Hospital of USC | 2 |
| COUC-TX1 University of Colorado Hospital/Health Sc | 1 |
| CTHH-TX1 Hartford Hospital | 3 |
| CTYN-TX1 Yale New Haven Hospital | 4 |
| DCGU-TX1 Medstar Georgetown Transplant Institute | 1 |
| DCWR-TX1 Walter Reed National Military Medical Cen | 1 |
| FLCC-TX1 Cleveland Clinic Florida Weston | 2 |
| FLFH-TX1 AdventHealth Orlando | 2 |
| FLJM-TX1 Jackson Memorial Hospital University of M | 3 |
| FLMR-TX1 Memorial Regional Hospital | 1 |
| FLSL-TX1 Mayo Clinic Hospital Florida | 1 |
| FLTG-TX1 Tampa General Hospital | 1 |
| GAEM-TX1 Emory University Hospital | 2 |
| IAIV-TX1 University of Iowa Hospitals and Clinics | 1 |
| IAVA-TX1 The Iowa City VA Health Care System | 1 |
| ILLU-TX1 Loyola University Medical Center | 2 |
| ILMM-TX1 Springfield Memorial Hospital | 1 |
| ILNM-TX1 Northwestern Memorial Hospital | 2 |
| ILPL-TX1 Rush University Medical Center | 1 |
| ILSF-TX1 OSF Saint Francis Medical Center | 1 |
| INIM-TX1 Indiana University Health | 1 |

| | |
|---|---|
| INSV-TX1 Ascension St. Vincent Hospital | 1 |
| KSUK-TX1 University of Kansas Hospital | 1 |
| MAMG-TX1 Massachusetts General Hospital | 1 |
| MAUM-TX1 UMass Memorial Medical Center | 1 |
| MDJH-TX1 Johns Hopkins Hospital | 8 |
| MDUM-TX1 University of Maryland Medical System | 6 |
| MIBH-TX1 William Beaumont Hospital | 1 |
| MIHF-TX1 Henry Ford Hospital | 1 |
| MIUM-TX1 University of Michigan Medical Center | 1 |
| MORH-TX1 Research Medical Center | 2 |
| MSUM-TX1 University of Mississippi Medical Center | 1 |
| NCBG-TX1 Wake Forest Baptist Medical Center | 8 |
| NCCM-TX1 Carolinas Medical Center | 1 |
| NEUN-TX1 The Nebraska Medical Center | 1 |
| NJHK-TX1 Hackensack University Medical Center | 1 |
| NJRW-TX1 Robert Wood Johnson University Hospital | 3 |
| NJSB-TX1 Saint Barnabas Medical Center | 3 |
| NYAM-TX1 Albany Medical Center Hospital | 1 |
| NYCC-TX1 Long Island Jewish Medical Center-Cohen C | 1 |
| NYCP-TX1 NY Presbyterian Hospital/Columbia Univ. M | 404 |
| NYDS-TX1 State University of New York, Downstate M | 76 |
| NYEC-TX1 Erie County Medical Center | 13 |
| NYFL-TX1 Strong Memorial Hospital, University of R | 4 |
| NYMA-TX1 Montefiore Medical Center | 408 |
| NYMS-TX1 Mount Sinai Medical Center | 253 |
| NYNS-TX1 North Shore University Hospital/Northwell | 59 |
| NYNY-TX1 New York-Presbyterian Hospital/Weill Corn | 233 |
| NYSB-TX1 University Hospital of State University o | 191 |
| NYSL-TX1 St. Luke's Roosevelt Hospital Center | 7 |
| NYUC-TX1 NYU Langone Health | 144 |
| NYUM-TX1 State University of New York Upstate Medi | 4 |
| NYVA-TX1 James J. Peters VA Medical Center | 5 |
| NYWC-TX1 Westchester Medical Center | 84 |
| OHCO-TX1 University of Toledo Medical Center | 2 |
| OHMV-TX1 Miami Valley Hospital | 1 |
| OHTC-TX1 The Christ Hospital | 1 |
| PAHH-TX1 Pinnacle Health System at Harrisburg Hosp | 1 |
| PALH-TX1 The Lankenau Hospital | 2 |
| PAPT-TX1 University of Pittsburgh Medical Center | 1 |

| | |
|---|---|
| PATJ-TX1 Thomas Jefferson University Hospital | 1 |
| PAUP-TX1 Hospital of the University of Pennsylvani | 2 |
| TNEM-TX1 Erlanger Medical Center | 1 |
| TNST-TX1 Saint Thomas Hospital | 1 |
| TNUK-TX1 University of Tennessee Medical Center at | 2 |
| TNVU-TX1 Vanderbilt University Medical Center | 1 |
| TXHH-TX1 Memorial Hermann Hospital, University of | 1 |
| TXHI-TX1 CHI St. Luke's Health Baylor College of M | 1 |
| TXHS-TX1 Methodist Specialty and Transplant Hospit | 2 |
| TXJS-TX1 University of Texas Medical Branch at Gal | 2 |
| TXMC-TX1 Methodist Dallas Medical Center | 1 |
| TXSP-TX1 UT Southwestern Medical Center/William P. | 1 |
| TXSW-TX1 Scott and White Memorial Hospital | 1 |
| TXTX-TX1 Baylor University Medical Center | 1 |
| VAFH-TX1 Inova Fairfax Hospital | 2 |
| VAMC-TX1 VCU Health System Authority, VCUMC | 1 |
| VTMC-TX1 The University of Vermont Medical Center | 1 |
| WASH-TX1 Providence Sacred Heart Medical Center & | 1 |
| WAVM-TX1 Virginia Mason Medical Center | 1 |
| WIUW-TX1 University of Wisconsin Hospital and Clin | 1 |

**Table S2.** Accuracy summary of kidney tissue compartment prediction model on independent testing set images.

| Group | TPR | PPV | F-score |
|---|---|---|---|
| Normal Glomerulus | 0.96 | 0.97 | 0.97 |
| Sclerotic Glomerulus | 0.91 | 0.93 | 0.92 |
| Artery | 0.94 | 0.91 | 0.94 |
| Arterial Intimal Thickening | 0.94 | 0.92 | 0.93 |
| All Tubule | 0.90 | 0.87 | 0.90 |

**Table S3.** Correlation of digital scores and pathologist scores with 6-month, 12-month, and 24-month eGFR post-transplant after adjusting for recipient and peri-transplant information.

| Outcome | Scores | P-value | Coefficient | P-value | Coefficient |
|---|---|---|---|---|---|
| | | By Machine | | By Pathologist | |
| **M6 eGFR** | Sclerotic Glomeruli% | 2.58E-08 | -60.14 | 1.28E-04 | -6.60 |
| | Arterial Intimal Fibrosis% | 1.09E-13 | -31.48 | 4.63E-11 | -6.59 |
| | Interstitial Fibrosis% | 1.97E-04 | -16.99 | 4.16E-03 | -4.50 |
| **M12 eGFR** | Sclerotic Glomeruli% | 5.75E-11 | -74.69 | 1.12E-03 | -6.06 |
| | Arterial Intimal Fibrosis% | 2.33E-12 | -31.68 | 5.91E-13 | -7.69 |
| | Interstitial Fibrosis% | 7.25E-05 | -19.23 | 2.69E-03 | -5.02 |
| **M24 eGFR** | Sclerotic Glomeruli% | 2.48E-10 | -104.61 | 4.63E-03 | -7.13 |
| | Arterial Intimal Fibrosis% | 8.06E-10 | -38.59 | 4.87E-08 | -8.09 |
| | Interstitial Fibrosis% | 3.35E-04 | -23.21 | 1.97E-02 | -5.62 |

**Table S4.** Demographic and clinical characteristics of recipients and donors between Survival Discovery Set and Validation Set.

| Characteristics | Discovery (n=520) | Validation (n=1040) | P-value |
|---|---|---|---|
| Recipient Age | 54.37 ±13.55 | 55.08 ±13.15 | 0.322 |
| Recipient Gender | | | 0.58 |
| Female | 200(38.46) | 385(37.02) | |
| Male | 320(61.54) | 655(62.98) | |
| Recipient Race | | | 0.478 |
| Asian | 61(11.73) | 112(10.77) | |
| African American | 187(35.96) | 421(40.48) | |
| Hispanic | 138(26.54) | 249(23.94) | |
| Caucasian | 130(25) | 252(24.23) | |
| Other | 4(0.77) | 6(0.58) | |
| Recipient Pre-transplant Dialysis | | | 0.114 |
| No | 64(12.36) | 100(9.62) | |
| Yes | 454(87.64) | 939(90.38) | |
| Estimated Post-Transplant Survival (EPTS) score | 0.54±0.29 | 0.55±0.3 | 0.724 |
| Pump Usage | | | 0.411 |
| No | 42(8.08) | 72(6.92) | |
| Yes | 478(91.92) | 968(93.08) | |
| Cold Ischemia Time (hours) | 25.73 ±10.78 | 26.72 ±11.14 | 0.093 |
| Induction Type | | | 0.166 |
| Lymphocyte Non-depletion | 103(19.81) | 205(19.71) | |
| Lymphocyte Depletion | 389(74.81) | 752(72.31) | |
| None | 28(5.38) | 83(7.98) | |
| Death Censored Graft Loss | | | 0.946 |
| No | 421(80.96) | 840(80.77) | |
| Yes | 99(19.04) | 200(19.23) | |
| Follow-up Days | 1929.48 ±952.88 | 1959.18 ±957.9 | 0.563 |
| Donor Age | 45.22 ±14.13 | 44.97 ±13.25 | 0.75 |
| Donor Gender | | | 0.522 |
| Female | 187(39.62) | 357(41.56) | |
| Male | 285(60.38) | 502(58.44) | |
| Donor Race | | | 0.98 |
| Asian | 16(3.39) | 24(2.79) | |
| African American | 73(15.47) | 137(15.95) | |
| Hispanic | 96(20.34) | 174(20.26) | |
| Caucasian | 283(59.96) | 516(60.07) | |
| Other | 4(0.84) | 8(0.93) | |
| KDPI | 0.54 ±0.26 | 0.54 ±0.25 | 0.99 |

* P-values are calculated by Fisher's exact test (categorical variables) or Student's t-test (continuous variables).

**Table S5.** Association of digital scores and pathologist scores with graft loss in Survival Discovery Set (n=520) after adjusting for recipient and peri-transplant information.

| **Scores** | | **PH Assumption P-value** | **Coefficient P-value** | **Hazard Ratio** | **Lower 95% CI** | **Upper 95% CI** |
|---|---|---|---|---|---|---|
| Sclerotic Glomeruli% | by machine | 8.83E-01 | 3.56E-03 | 496.39 | 7.64 | 32271.11 |
| | by pathologists | 2.55E-01 | 2.97E-01 | 1.38 | 0.75 | 2.55 |
| Arterial Intimal Fibrosis% | by machine | 9.32E-01 | 2.30E-02 | 6.11 | 1.28 | 29.08 |
| | by pathologists | 3.55E-01 | 4.13E-01 | 1.19 | 0.78 | 1.81 |
| Interstitial Fibrosis% | by machine | 2.31E-01 | 2.60E-02 | 5.53 | 1.23 | 24.89 |
| | by pathologists | 9.30E-01 | 1.66E-01 | 1.63 | 0.82 | 3.24 |

*Cox p-values are calculated by Wald test from Cox proportional hazards regression. The proportional hazards assumptions are assessed through chi-square goodness of fit test between Schoenfeld residuals and time. Non-significant p-values confirm the assumption.

**Table S6.** Association of combined digital scores and KDPI with graft loss in Survival Discovery Set (n=520) after adjusting for recipient and peri-transplant information.

| Scores | PH Assumption P-value | Coefficient P-value | Hazard Ratio | Lower 95% CI | Upper 95% CI |
|---|---|---|---|---|---|
| Combined Digital Score | 3.50E-01 | 1.92E-05 | 1.43 | 1.21 | 1.68 |
| KDPI | 8.61E-01 | 1.21E-02 | 3.17 | 1.29 | 7.83 |

*Cox p-values are calculated by Wald test from Cox proportional hazards regression. The proportional hazards assumptions are assessed through chi-square goodness of fit test between Schoenfeld residuals and time. Non-significant p-values confirm the assumption.

**Table S7.** Association of Kidney Donor Quality Score and other public composite pathological scores with graft loss in Survival Discovery Set (n=520) after adjusting for recipient and peri-transplant information.

| Scores | PH Assumption P-value | Coefficient P-value | Hazard Ratio | Lower 95% CI | Upper 95% CI |
|---|---|---|---|---|---|
| Kidney Donor Quality Score | 2.55E-01 | 5.62E-06 | 1.34 | 1.18 | 1.52 |
| Chronic Damage Score | 8.56E-01 | 1.48E-01 | 1.22 | 0.93 | 1.61 |
| Leuven Score | 8.01E-01 | 7.89E-02 | 1.01 | 1.00 | 1.02 |

*Cox p-values are calculated by Wald test from Cox proportional hazards regression. The proportional hazards assumptions are assessed through chi-square goodness of fit test between Schoenfeld residuals and time. Non-significant p-values confirm the assumption.

**Table S8.** Coefficients of 1-year and 4-year graft loss prediction logistic regression models by incorporating clinical factors and Kidney Donor Quality Score.

**1-YEAR MODEL**

| | **Estimate** | **Lower 95% CI** | **Upper 95% CI** |
|---|---|---|---|
| (Intercept) | -1.22 | -3.06 | 0.49 |
| Cold Ischemia Time (hours) | 0.03 | -0.01 | 0.07 |
| Use of Pump (Y) | -1.56 | -2.65 | -0.34 |
| Induction Type (Lymphocyte Depletion) | -2.70 | -3.86 | -1.52 |
| Induction Type (Lymphocyte Non-depletion) | -2.09 | -3.45 | -0.79 |
| Kidney Donor Quality Score | 0.37 | 0.12 | 0.63 |

**4-YEAR MODEL**

| | **Estimate** | **Lower 95% CI** | **Upper 95% CI** |
|---|---|---|---|
| (Intercept) | 0.12 | -1.46 | 1.64 |
| Cold Ischemia Time (hours) | 0.02 | -0.01 | 0.05 |
| Recipient Age | -0.04 | -0.06 | -0.02 |
| Induction Type (Lymphocyte Depletion) | -1.76 | -2.75 | -0.73 |
| Induction Type (Lymphocyte Non-depletion) | -1.88 | -3.06 | -0.71 |
| Kidney Donor Quality Score | 0.40 | 0.22 | 0.59 |

**Table S9.** Graft outcomes within 1-year post-transplant of transplanted kidneys identified as extreme high risk and should have been considered for discard by KDQS or 1-year survival prediction model. The 1-year risk was defined as: 1) graft loss at 1-year, or 2) severe loss of kidney function at last follow-up within 1-year.

| ID | Identified By | KDQS | Prediction Score By 1-Year Model | M6 eGFR | M12 eGFR | Follow-Up Days | Graft Loss | Recipient Death | Graft Loss at M12 | 1-year Risk |
|---|---|---|---|---|---|---|---|---|---|---|
| **1** | KDQS | 8 | | 15.95 | | 284 | Yes | Yes | Yes | Yes |
| **2** | KDQS | 7 | | | | 0 | Yes | Yes | Yes | Yes |
| **3** | KDQS | 7 | | | | 0 | Yes | Yes | Yes | Yes |
| **4** | KDQS | 7 | | 63.88 | 87.28 | 336 | No | Yes | No | No |
| **5** | KDQS | 8 | | 31.45 | 30.24 | 341 | No | No | | Yes |
| **6** | KDQS | 8 | | 28.92 | 20.92 | 531 | No | No | No | Yes |
| **7** | KDQS | 7 | | 41.74 | 31.19 | 358 | No | No | | Yes |
| **8** | KDQS | 7 | | 17.75 | 15.24 | 648 | Yes | Yes | No | Yes |
| **9** | KDQS | 7 | | 27.13 | 24.74 | 665 | Yes | Yes | No | Yes |
| **10** | 1-Year Model | 6 | 0.62 | 36.78 | | 336 | Yes | Yes | Yes | Yes |
| **11** | 1-Year Model | 2 | 0.61 | | | 126 | Yes | Yes | Yes | Yes |
| **12** | 1-Year Model | 6 | 0.59 | | | 101 | Yes | Yes | Yes | Yes |
| **13** | 1-Year Model | 2 | 0.68 | 55.77 | 22.49 | 2380 | No | No | No | Yes |
